# Nuclear–Cytoskeletal Uncoupling in Aging Capillaries Disrupts Endothelial Nuclear Flattening and Lumen Formation

**DOI:** 10.64898/2026.03.09.710624

**Authors:** Caitlin Francis, Felipe Leser, Roberto Lopez, Anne Eichmann

## Abstract

Loss of small blood vessels compromises tissue perfusion during aging, but the underlying mechanisms remain unclear. We investigated microvascular integrity in physiologically aged mice and in Hutchinson-Gilford Progeria Syndrome, a genetic disorder of premature vascular aging. Progeric and aged mice exhibited capillary loss and impaired perfusion, caused by cell autonomous defects in endothelial nuclei that failed to flatten and instead obstructed vessel lumens. Progerin expression in endothelial cells altered LINC complex composition by increasing SUN proteins and reducing nuclear-envelope Nesprin engagement, disrupting coupling to cytoskeletal actin and vimentin. Experimental disruption of nuclear-cytoskeletal coupling impaired nuclear flattening and lumen formation, whereas restoration improved nuclear morphology and vessel formation in progeric cells. Aged mouse and human endothelium exhibited changes in LINC complex and vimentin expression. Pharmacological Progerin inhibition restored nuclear-cytoskeletal coupling and lumen formation in patient-derived endothelial cells. These findings implicate defective nuclear-cytoskeletal coupling in microvascular dropout in Progeria and physiological aging.

## Introduction

Microvascular functional decline is a hallmark of vascular aging that compromises tissue perfusion, yet the cellular mechanisms driving capillary loss remain poorly understood. Efficient oxygen and nutrient exchange depend on a specialized endothelial monolayer that can be as thin as 200 nm, these dimensions impose unique morphological constraints requiring endothelial nuclei to flatten and reposition to maintain vessel patency.^1,29,33^ How endothelial nuclei undergo these morphological changes, and whether their disruption contributes to microvascular dysfunction, remain poorly understood.

Hutchinson–Gilford Progeria Syndrome (hereafter Progeria) is a rare, fatal genetic disorder caused by the accumulation of Progerin, a mutant Lamin-A isoform at the inner nuclear membrane that disrupts nuclear structure and function.^3,13,15,16^ Children with Progeria develop severe cardiovascular pathology that shares features with age-associated vascular disease.^2,20,30,35^ Previous studies have established profound vascular smooth muscle cell loss and macrovascular pathology in Progeria, but the effects of Progerin on the microvasculature remain largely unexplored.^21,26,36^

Nuclear positioning and morphology are influenced by mechanical interactions between the nucleus and cytoskeleton, mediated in part by the Linker of Nucleoskeleton and Cytoskeleton (LINC) complex.^5,12,27^ The LINC complex spans the nuclear envelope through interactions between Lamin-A and SUN (Sad1 and UNC-84 homology) proteins in the inner nuclear membrane and SUN and Nesprins (nuclear envelope spectrin repeat proteins) in the outer nuclear membrane.^4,19,46^ Nesprin-1 and -2 connect the nucleus to actin filaments, whereas Nesprin-3 associates with vimentin intermediate filaments through plectin, enabling cytoskeletal forces to be transmitted to the nuclear lamina.^7,25,28^ Previous studies have demonstrated that forces transmitted through the LINC complex protein Nesprin-1 regulate endothelial adhesion, migration, and alignment in response to mechanical cues while preserving chromatin organization and nuclear integrity.^6,11^ Consistent with these findings, multiple reports have shown that LINC complex proteins are required for endothelial adaptation to shear stress, cyclic stretch, and other hemodynamic forces.^31,42,44^ However, whether LINC-mediated nuclear-cytoskeletal coupling regulates endothelial nuclear flattening and positioning during capillary morphogenesis, and whether disruption of this interface contributes to microvascular dysfunction, remains unknown.

Here, we identify microvascular dropout and impaired perfusion in progeric mice, accompanied by endothelial nuclear abnormalities that obstruct capillary lumens. Using complementary mouse and human endothelial models, we demonstrate that Progerin disrupts nuclear positioning, cytoskeletal organization, and vessel morphogenesis through alterations in LINC complex composition. Experimental disruption of nuclear-cytoskeletal coupling partially reproduces these defects, supporting its functional contribution to endothelial morphogenesis. We further identify abnormalities in endothelial nuclear positioning, perinuclear vimentin organization, and LINC complex components during physiological aging, revealing vulnerabilities of the nuclear-cytoskeletal interface in both conditions. Finally, pharmacological inhibition of Progerin restores LINC-associated protein organization and improves vessel formation in patient-derived endothelial cells. Together, these findings identify nuclear–cytoskeletal coupling as an important regulator of endothelial morphogenesis and implicate its disruption in microvascular pathology.

## Results

### Progeric and Aged Mice Exhibit Microvascular Dropout

We analyzed mice expressing human Progerin (LMNA G608G), which recapitulate key features of human disease and exhibit a reduced lifespan of approximately 25 weeks in our facility. By 20 weeks of age, homozygous LMNA G608G mice, hereafter referred to as progeric mice, exhibited advanced progeroid pathology, akin to human patients, accompanied by a rapid decline in overall health, while their wildtype littermate controls remained healthy. We assessed microvascular density across gut, brain, and retina in 20-week-old progeric mice, 20-week-old wildtype littermate controls and 2-year-old aged wild-type mice, using an antibody recognizing the endothelial luminal marker podocalyxin (Podxl). Compared to controls, both progeric and aged mice exhibited significantly reduced vessel density across the tissues examined (**Figure 1a-d**). To further characterize these microvascular abnormalities, we stained retinal whole mounts for Podxl and ETS-related gene (ERG), an endothelial nuclear marker. Lumen narrowing was significantly increased in both progeric and aged mice compared to controls (**Figure 1e-f**) and coincided with the presence of an endothelial nucleus obstructing the lumen in a greater proportion of cases in progeric and aged mice, with increases of 24% and 16%, respectively, compared to controls (**Figure 1g**). Analysis of 10-week-old progeric mice revealed no quantifiable microvascular defects compared to controls (**Supplemental Figure 1a-d**), indicating that microvascular dropout occurred between 10 and 20 weeks of age.

**Figure 1.**
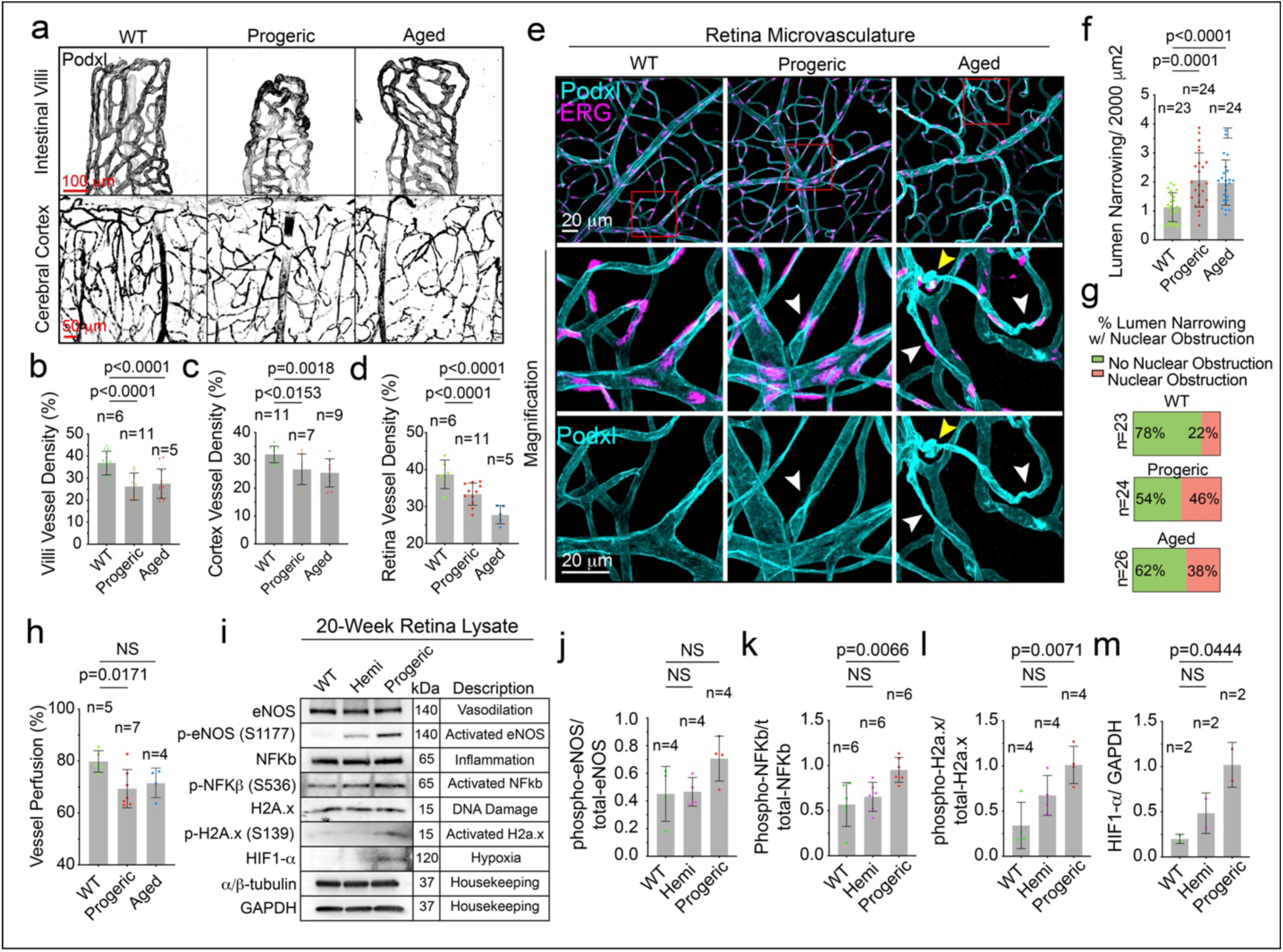
Progeric and Aged Mice Exhibit Microvascular Dropout. a) Image representatives of microvasculature, marked by podocalyxin (podxl), of the intestinal villi and cerebral cortex. Quantification of b) villi vessel density, c) cortex vessel density and d) retina vessel density. e) Image representatives of retina vasculature marked by podxl and endothelial nuclear marker, ERG. White arrowheads denote lumen narrowing by nuclear obstruction and yellow arrowheads denote vascular tortuosity. Red box denotes area of magnification. Quantification of f) instances of lumen narrowing by area, g) percent lumen narrowing with nuclear obstruction, and h) percent vessel perfusion with Sulfo-NHS-Biotin. i) Western blotting of whole retina lysate for indicated proteins. Quantification of j) serine-1177 phosphorylated-eNOS by total eNOS, k) serine-536 phosphorylated NFKb by total NFKb, l) serine-139 phosphorylated H2A.x by total H2A.x and m) total HIF-1α by total GAPDH. N= number of separate lysates. In all quantification a minimum of 4 biological replicates were used per condition.

To assess the extent of capillary blood flow, mice were systemically perfused with Sulfo-NHS-Biotin. Progeric and aged mice exhibited significantly reduced capillary perfusion compared with age-matched controls (**Figure 1h**). To investigate the molecular consequences associated with reduced perfusion, we isolated whole retinas and analyzed them by western blotting (**Figure 1f-j**). Levels of hypoxia-inducible factor-1α (HIF-1α) were elevated in homozygous progeric retinas, indicating increased tissue hypoxia. We next assessed endothelial nitric oxide synthase (eNOS) signaling by measuring total eNOS and phosphorylated eNOS (Ser1177). Progeric retinas exhibited markedly increased phospho-eNOS levels, suggesting activation of compensatory vasodilatory pathways in response to vascular dysfunction. Finally, we examined markers of DNA damage and inflammation, including H2A.x and NF-κB. Progeric mice showed significant increases in both markers, consistent with elevated DNA damage and inflammatory signaling **Figure 1f-j**). Hemizygous progeric mice showed no significant changes in signaling proteins or vascular defects at 20-weeks (**Figure 1f-j** and data not shown), we therefore focused on the homozygous progeric mice.

Together, these findings demonstrate that Progerin expression causes progressive microvascular dysfunction characterized by impaired tissue perfusion, vessel rarefaction, endothelial lumen obstruction, and activation of hypoxic and stress-response pathways. The presence of similar microvascular abnormalities in aged wild-type mice suggests that these defects may reflect conserved mechanisms of vascular decline, prompting us to query whether they were endothelial cell autonomous or imposed by the environment.

### Progerin-Induced Endothelial Defects are Cell Autonomous

To determine whether the vascular abnormalities observed in vivo reflect endothelial cell-autonomous defects, we isolated lung endothelial cells from progeric, aged, and wild-type mice and assessed sprouting angiogenesis using an ex vivo microcarrier bead assay (**Figure 2a**). This approach enables the evaluation of vessel formation independently of the surrounding tissue environment while retaining the genetic background and prior physiological exposure of donor cells. Sprouts derived from progeric and aged endothelial cells were significantly shorter than WT controls and exhibited a reduced lumen diameter (**Figure 2b–d**). Consistent with our in vivo observations, endothelial nuclei frequently occupied the central region of progeric and aged sprouts, obstructing the space normally occupied by the lumen (**Figure 2b,e, white arrowheads**). Nuclear width was significantly increased in both groups, accompanied by a 16% and 22% increase in nuclear protrusion into the lumen in progeric and aged endothelial cells, respectively, compared to controls (**Figure 2f–g**). Together, these findings demonstrate that endothelial cells from progeric and aged mice retain defects in nuclear positioning, sprout extension, and lumen formation outside their native tissue environment.

**Figure 2.**
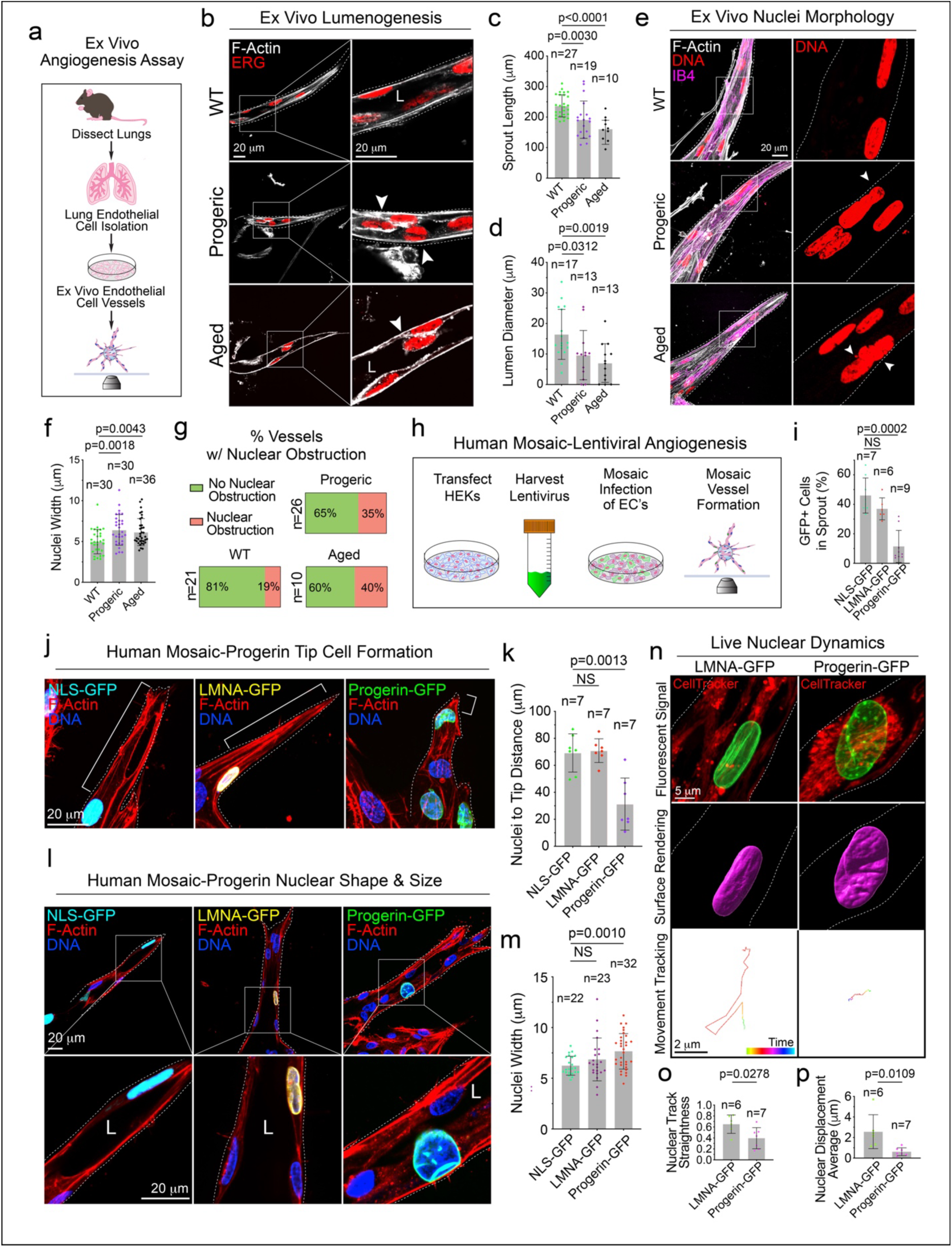
Sprouting Assays Reveal Conserved Endothelial Cell Specific Abnormalities in Mouse and Human. a) Cartoon outline of experimental procedure for ex vivo angiogenesis assay. b) 60x z-slice image representatives of vessel lumen formation. White arrowheads denote abnormally localized nuclei. Quantification of c) sprout length and d) lumen diameter measured at the location of cell nuclei within the vessel shaft. e) 60x z-projection image representatives of nuclear morphology and clustering. White arrowheads denote misshapen nuclei. Quantification of f) nuclear width and g) percentage of vessels with nuclei obstructing the lumen. h) Cartoon outline of experimental procedure for human (HUVEC) mosaic lentiviral infection angiogenesis assay. i) Quantification of GFP+ cells per sprout by total GFP+ cells on bead and in sprouts. N=number of beads. j) Z-projection image representatives of indicated condition tip cells. Bracket denotes measurement. k) Quantification of nuclei to tip distance. l) Z-slice image representatives of indicated conditions nuclear morphology. Box denotes area of magnification, L denotes lumen. m) Quantification of nuclear width. n) Z-projection stills of live imaged nuclei. Middle panels are nuclei surface rendering and bottom panels are of live imaging tracks. Quantification of o) nuclear track straightness and p) nuclear displacement average. N=number of nuclei live imaged. In all panels dashed line denotes vessel outline.

To assess whether Progerin was sufficient to induce these abnormalities in human endothelial cells, we transduced human umbilical vein endothelial cells (HUVECs) with lentiviral vectors encoding fluorescently tagged Progerin (**Figure 2h**). Vectors encoding nuclear localization sequence (NLS)-GFP or full-length Lamin-A (LMNA)-GFP served as controls. Viral titers were adjusted to achieve mosaic expression, enabling direct comparison of Progerin-expressing and non-expressing endothelial cells within the same sprout. We quantified sprout participation as the proportion of GFP-positive cells located within sprouts relative to the total GFP-positive population on the bead and within sprouts. Whereas more than 35% of NLS-GFP- or LMNA-GFP-expressing cells participated in sprout formation, only 10% of Progerin-GFP-positive cells entered sprouts (**Figure 2i**). Progerin-expressing cells were also rarely observed at the sprout tip. When they occupied this position, their nuclei were displaced toward the leading edge compared with control nuclei (**Figure 2j–k**). To determine whether impaired sprout extension was also evident in vivo, we examined the developing retinal vasculature of postnatal day 6 (P6) mice. Progeric retinas exhibited significantly reduced vascular outgrowth and shorter endothelial projections at the angiogenic front compared with WT controls, (**Supplemental Figure 2a–d**) demonstrating reduced growth at early stages that normalized at 10 weeks of age (see Supplemental Figure1a-d). These findings support a role for Progerin in disrupting endothelial sprout extension across experimental models.

Consistent with the primary mouse endothelial cell phenotype, Progerin-GFP-expressing HUVECs exhibited significantly increased nuclear width compared with NLS-GFP and LMNA-GFP controls (**Figure 2l–m**). Protrusion of Progerin-expressing nuclei into the lumen was accompanied by reduced lumen formation (**Supplemental Figure 2e-f**). Lumen diameter, measured at the location of the GFP-positive nuclei in the vessel stalk, was significantly smaller for Progerin-GFP vessels relative to both NLS-GFP and LMNA-GFP transduced control vessels. (**Supplemental Figure 2g**). To investigate whether these morphological abnormalities were associated with altered nuclear movement, we performed four-hour live-cell imaging of LMNA-GFP- and Progerin-GFP-expressing endothelial cells. Progerin-expressing nuclei displayed frequent local oscillations and reduced directional persistence compared with LMNA-expressing nuclei (**Supplemental Movies 1 and 2**). Quantitative tracking revealed significant differences in nuclear track straightness and average displacement between Progerin- and LMNA-expressing cells (**Figure 2n–p**), demonstrating that Progerin alters endothelial nuclear dynamics during sprouting.

Because directional nuclear movement is closely associated with cell polarity and migration, we next investigated whether Progerin disrupts endothelial polarization. In a migration assay, Progerin-expressing cells exhibited impaired Golgi reorientation toward the direction of migration and reduced migration distances (**Supplemental Figure 2h–j**). Similarly, Golgi positioning was disrupted in three-dimensional progeric endothelial sprouts, where Golgi apparatuses exhibited poor orientation relative to the sprout tip and frequently appeared fragmented (**Supplemental Figure 2k-m**). Together, these findings demonstrate that Progerin expression is associated with impaired nuclear dynamics, cellular polarity, and directional migration, providing a mechanistic basis for defective endothelial sprouting. Importantly, endothelial cells isolated from aged mice exhibited similar abnormalities in nuclear positioning and sprout formation, identifying a shared cellular phenotype between Progeria and physiological aging.

### Cytoskeletal Defects in the Aortic Endothelium of Aged and Progeric Mice

Impaired nuclear morphology and positioning may reflect defects in cytoskeletal function, particularly in the mechanisms that mediate coupling between the nucleus and the cytoskeleton at the nuclear-cytoskeletal interface. To understand how Progerin impacts this interface, we imaged the cytoskeleton of human vessels transduced with Progerin-GFP or control LMNA-GFP, as described above. We examined filamentous (f-) actin and the intermediate filament vimentin. Vimentin forms a key structural link with the nucleus and mutations in this protein have caused pre-mature aging phenotypes.^10^ Because vimentin antibodies did not give us high-quality immunostaining in 3-D, we probed for vimentin by co-transducing with vimentin-mCherry and for f-actin by immunolabeling. Both f-actin and vimentin localized abnormally in Progerin-GFP+ nuclei compared with control LMNA-GFP+ nuclei, with significantly reduced perinuclear localization that affected vimentin most severely (**Figure 3a-c**).

**Figure 3.**
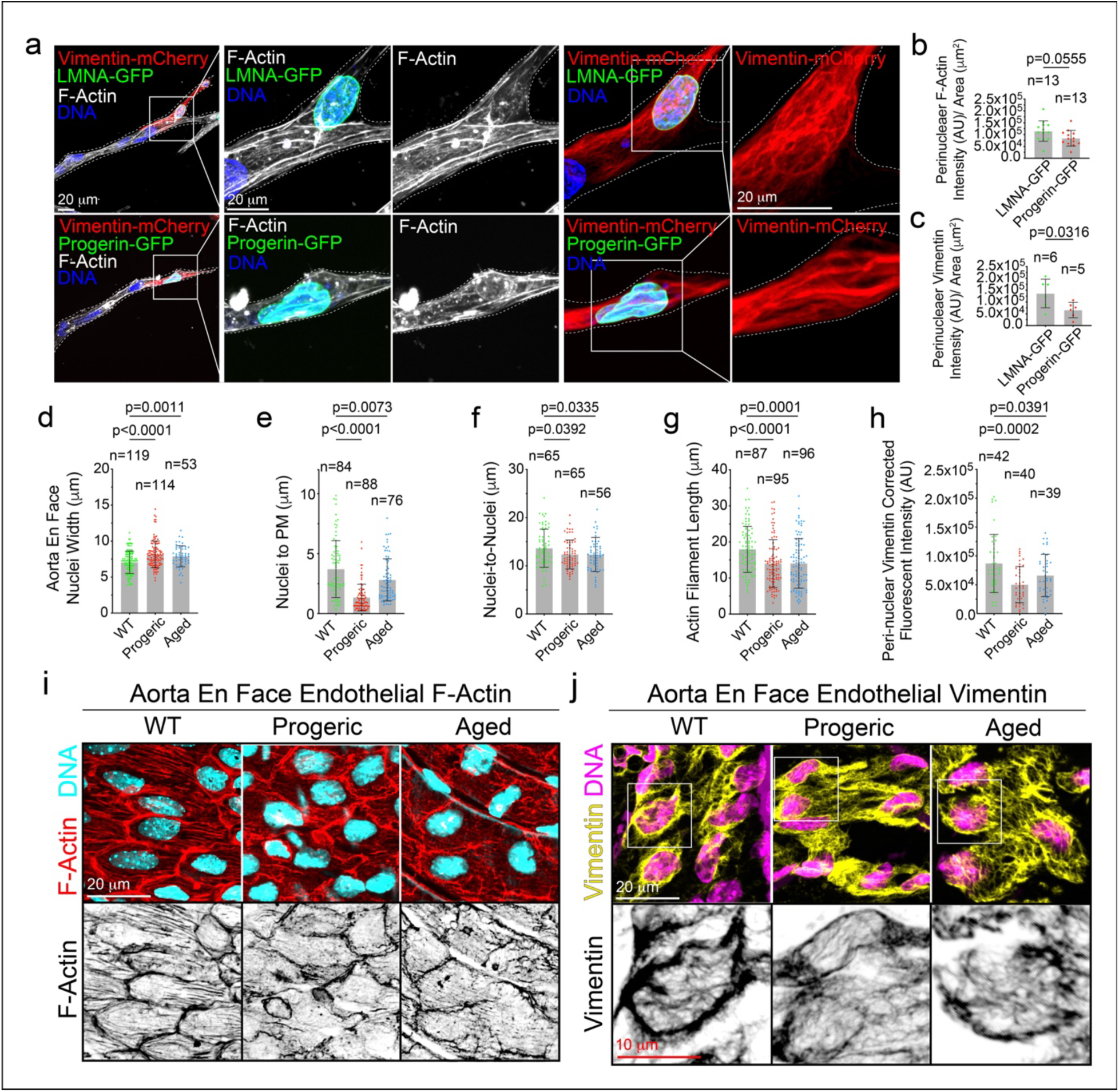
Progeric Aged- and 20-week Aortic Endothelial Cells have Cytoskeletal Defects. a) 60x z-projection image representatives of human in vitro grown vessels with Lamin-A (top panels) and progerin (bottom panels) mosaic lentiviral infection with f-actin stain and vimentin-mCherry transduction. Quantification of b) perinuclear fluorescent intensity of f-actin and c) perinuclear fluorescent intensity of vimentin-mCherry. Quantification of d) nuclear width, e) nuclei to plasma membrane distance, f) nuclei to nuclei distance, g) actin filament length and h) nuclear vimentin fluorescent intensity. All nuclei were quantified within inner aortic arch region. i) Image representatives of aorta en face inner curvature stained for f-actin (filamentous actin) and DNA. j) Image representatives of aorta en face inner curvature stained for vimentin and DNA. Box denotes area of magnification. For all quantification a minimum of 3 biological replicates were measured.

To determine if these findings are consistent in vivo and in aged endothelium, we performed en face imaging of aortic endothelial cells, which enables high-resolution visualization of endothelial morphology, protein localization, and subcellular organization within the intact vessel wall. Nuclear width was significantly greater in progeric and aged endothelial cells compared to 20-week-old WT controls (**Figure 3d, i**) and nuclear positioning was different. Progeric and aged endothelial nuclei were more likely to make contact with the plasma membrane or near neighboring nuclei rather than centralized within the cell as WT nuclei were (**Figure 3e-f, i**).

Staining for f-actin revealed reduced cortical actin localization and significantly shorter filament lengths in endothelial cells from progeric mice as well as aged mice compared with wildtype controls (**Figure 3g,i**). Vimentin staining revealed a marked reduction in perinuclear vimentin density in both progeric and aged endothelial cells compared with controls (**Figure 3h, j**). In addition to reduced abundance around the nucleus, vimentin organization was visibly disrupted in progeric and aged endothelial cells, suggesting compromised nuclear-cytoskeletal coupling.

Importantly, these endothelial cytoskeletal abnormalities were evident before the onset of significant SMC loss in progeric mice, as our analysis of 10-week progeric mice revealed significant differences in f-actin and vimentin where smooth muscle cell coverage was intact. These data indicate that endothelial nuclear-cytoskeletal dysfunction is likely a primary event rather than a secondary consequence of vascular remodeling (**Supplemental Figure 3a-i**). This early aortic involvement is particularly relevant given the severe aortic pathology documented in children with Progeria.^26,36^ Collectively, these findings identify nuclear-cytoskeletal interface disruption, involving both actin and vimentin networks, as an early feature of progeric vascular pathology and reveal parallels with endothelial aging. We next set out to understand the mechanisms underlying nuclear-cytoskeletal uncoupling.

### Aging and Progerin Impairs the LINC Complex in Endothelial Cells

The loss of perinuclear cytoskeletal protein localization in progeric endothelial cells suggested potential disruption of the LINC complex. To investigate alterations in LINC complex components (**Figure 4a**), we first examined SUN1 and SUN2, both inner nuclear membrane proteins that bind Lamin-A and link the nuclear lamina to Nesprins. SUN1 expression was higher in Progerin-expressing cells than in controls (**Figure 4b-c**), consistent with previous reports.^8,9^ We also found increased SUN2 expression in Progerin-expressing cells compared with controls (**Figure 4b,d**).

**Figure 4.**
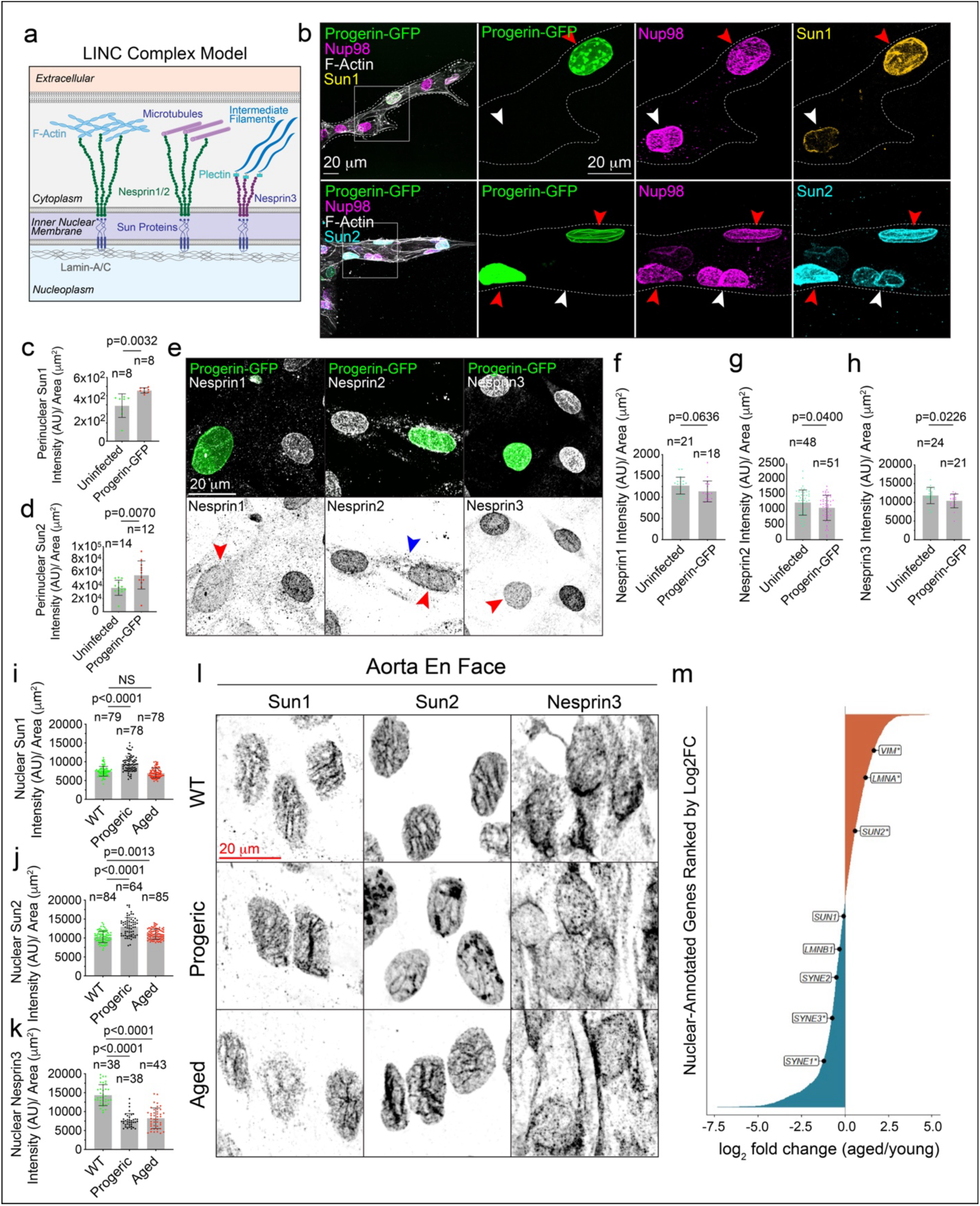
Endothelial Nuclear Membrane Properties are Altered by Progerin Expression. a) Cartoon model of LINC complex protein composition and organization. b) 60x z-projection image representatives of Progerin-GFP mosaic expression and staining for Sun1 (top row) and Sun2 (bottom row). Red arrowhead denotes Progerin+ cell and white arrowhead denotes Progerin-cell. Quantification of perinuclear c) Sun1 and d) Sun2 in 3-dimensional in vitro grown vessels. e) Image representatives of Lamin-A and Progerin expression with mCherry-Tubulin (microtubule marker). e) Image representatives of mosaic Progerin-GFP expression with Nesprin-1, Nesprin-2, and Nesprin-3 staining. Red arrowhead indicates cell expressing Progerin. Blue arrowhead denotes the cytoplasmic Nesprin-2 signal. Quantification of f) Nesprin-1, g) Nesprin-2, and h) Nesprin-3 fluorescent intensity by area in Progerin-GFP negative and positive nuclei. Quantification of nuclear i) Sun1 signal and j) Sun2 signal in aorta en face. k) Z-projections of Sun1 (first column), Sun2 (second column), and Nesprin3 (third column) in wildtype, progeric, and aged aorta en face. l) Quantification of nuclear Nesprin3 signal. m) Age-associated nuclear transcriptome remodeling in human heart endothelial cells. Ranked distribution of 5,328 nuclear-annotated genes (GO:0005634) in human heart ECs. Differential gene expression was assessed using donor-level pseudo-bulk RNA-sequencing analysis with limma–voom, comparing aged and young donors (n= 17.231 cells from 10 young patients and 81.907 cells from 12 aged participants), and ranked by its Log_2_FC. Selected components of the nuclear lamina–LINC–cytoskeleton axis (LMNA, LMNB1, SUN1, SUN2, SYNE1, SYNE2, SYNE3, and VIM) are highlighted. * Benjamini–Hochberg-adjusted FDR < 0.05, **FDR < 0.01, ***FDR < 0.001.

Nesprins1-3 are outer nuclear membrane proteins that form the cytoplasmic link of the LINC complex and include Nesprin-1, Nesprin-2, and Nesprin-3.^26^ Using mosaic infection with Progerin-GFP and antibody staining for Nesprins, we compared the nuclear envelope coverage of Nesprin-1, Nesprin-2, and Nesprin-3 in Progerin-expressing (Progerin^+^) cells and neighboring non-expressing (Progerin^−^) cells within the same vessels. All three Nesprins exhibited reduced coverage at the nuclear envelope in Progerin^+^ cells (**Figure 4e–h**). Furthermore, Nesprin signals were frequently detected in the cytoplasm rather than being restricted to the nuclear envelope, as shown for Nesprin-2 (**Figure 4e, blue arrowhead**). Together, these findings indicate that Progerin-expressing endothelial cells exhibit increased SUN protein coverage and reduced Nesprin coverage at the nuclear envelope.

To further validate these findings, we isolated nuclear and cytoplasmic fractions from endothelial cells grown in two-dimensional culture and assessed SUN1, SUN2, Nesprin-1, Nesprin-2, Nesprin-3, vimentin, β-actin, β-tubulin, plectin, GAPDH, and Nup98 by western blotting (**Supplemental Figure 4a**). Nup98, used as a nuclear marker for normalization, was detected predominantly in the nuclear fraction, whereas GAPDH was detected predominantly in the cytoplasmic fraction and used to normalize cytoplasmic protein levels. Consistent with our imaging results, SUN1 and SUN2 levels were significantly higher in nuclear fractions from Progerin-expressing cells than in those from controls (**Supplemental Figure 4b-c**). Nesprin-1, Nesprin-2, and Nesprin-3 were detected in both nuclear and cytoplasmic fractions, with Nesprin-1 showing a significant reduction and Nesprin-2 and Nesprin-3 also exhibiting reduced levels in Progerin-expressing cells compared to controls (**Supplemental Figure 4d-f**). All detectable isoforms of Nesprin-1 and -2 were quantified, although only bands sized 150-250kDa are displayed. No quantifiable differences were observed in β-actin or α/β-tubulin levels between the nuclear and cytoplasmic fractions (**Supplemental Figure 4g-h**). In contrast, vimentin levels were significantly lower in the nuclear fraction of Progerin-expressing cells than in controls, consistent with the loss of the perinuclear vimentin cage observed by imaging (**Supplemental Figure 4i**). These results further support the conclusion that Progerin expression alters the distribution of LINC complex components and associated cytoskeletal proteins.

Plectin, which links vimentin intermediate filaments to the nuclear envelope through Nesprin-3, was significantly reduced in both imaging analyses and nuclear fractions (**Supplemental Figure 4j-l**). ^34^ This finding further supports disruption of the vimentin-plectin-Nesprin-3 connection in Progerin-expressing cells. Collectively, our findings reveal an altered distribution of LINC complex components, characterized by increased SUN protein coverage and reduced Nesprin coverage at the nuclear envelope.

To determine if progeria and physiological aging were associated with alterations in LINC complex proteins in vivo, we used en face imaging of the aorta to examine SUN1, SUN2, and Nesprin-3 abundance in 20-week-old wild-type and progeric mice, as well as 2-year-old aged mice. Consistent with our in vitro findings, progeric mice exhibited increased SUN1 and SUN2 abundance and reduced Nesprin-3 abundance relative to controls (**Figure 4i–l**). Aged mice similarly exhibited increased SUN2 and reduced Nesprin-3, although SUN1 was not significantly different compared to controls.

To query whether associated genes are transcriptionally altered during physiological human aging, we analyzed the Human Aging Endothelial Cell Atlas (HAECA), a single-cell transcriptomic dataset encompassing 375,829 endothelial cells from 290 samples across 12 healthy human tissues and 34 publicly available single-cell scRNA-sequencing studies.14 In cardiac endothelial cells, aging was associated with significantly increased expression of *VIM* (vimentin) and *LMNA* (Lamin-A), elevated *SUN2*, unchanged *SUN1,* and reduced expression of *LMNB1* (Lamin-B1), *SYNE1* (Nesprin-1), *SYNE2* (Nesprin-2), and *SYNE3* (Nesprin-3) (**Figure 4m**). Notably, increased expression of *VIM* and *LMNA* occurred alongside reduced expression of Nesprin-encoding genes, revealing transcriptional changes in nuclear structural components and proteins that mediate nuclear-cytoskeletal connectivity. The resulting imbalance may affect the nuclear-cytoskeletal interface.

Across vascular beds in different tissues, age-associated changes in LINC complex gene expression were heterogeneous, with distinct expression patterns across tissues (**Supplemental Figure 4m).** Consistent across tissues, *SUN1* and *SYNE3* expression were relatively preserved in most endothelial populations, whereas *LMNA, VIM, SYNE1*, and *SUN2* exhibited more pronounced age-associated variation. Endothelial cells from kidney, breast, adipose, intestine, and muscle exhibited transcriptional patterns similar to the heart, while uterus, brain, skin, liver and lung endothelium displayed largely opposing patterns. In these latter tissues, aging was generally associated with reduced *VIM, LMNA*, and *SUN2* expression alongside increased expression of *SYNE1–3*. These findings suggest that aging differentially alters the relative expression of nuclear structural components and their cytoskeletal linkers across vascular beds.

Gene set enrichment analysis of nuclear architecture-related pathways resulted in aged cardiac endothelial cells exhibited significant positive enrichment of gene sets associated with nuclear membrane reassembly, nuclear envelope organization, and nuclear membrane organization, whereas these pathways did not reach statistical significance in most other endothelial populations (**Supplemental Figure 4n**). Given the pronounced cardiac pathology observed in children with Progeria, these findings suggest that LINC complex mediated nuclear-cytoskeletal coupling in cardiac endothelium is particularly susceptible to age-associated disruption.

Collectively, these findings demonstrate that Progerin expression alters the localization and abundance of LINC complex components, characterized by increased SUN protein levels, reduced Nesprin coverage at the nuclear envelope, and diminished association of cytoskeletal proteins with the nuclear compartment. Although physiological aging produces distinct molecular changes relative to Progeria, both conditions exhibit alterations in components of the nuclear– cytoskeletal interface. Together with our functional observations, these findings suggest that disruption of nuclear–cytoskeletal organization may contribute to age-associated endothelial nuclear abnormalities.

### LINC Complex Disruption Partially Phenocopies Progerin Expression

To test whether disruption of the LINC complex is central to the endothelial defects observed in progeric and aged endothelium, we expressed the dominant-negative KASH construct (mCherry-KASH-DN)^11^, which displaces endogenous Nesprins from the nuclear envelope and thereby disrupts nuclear-cytoskeletal coupling. Upon *in vitro* sprouting, HUVEC’s expressing mCherry-KASH-DN exhibited significantly abnormal nuclear morphology and blunted tip cells compared to control vessels (**Figure 5a–f**), demonstrating that disruption of LINC-mediated nuclear-cytoskeletal coupling is sufficient to induce several key features of the Progeria phenotype. However, when compared to effects of Progerin expression on lumen diameter (see Supplemental Figure 3c and Figure 5e), these defects were less severe, suggesting that additional mechanisms may contribute to dysfunction in Progerin-expressing endothelial cells.

**Figure 5.**
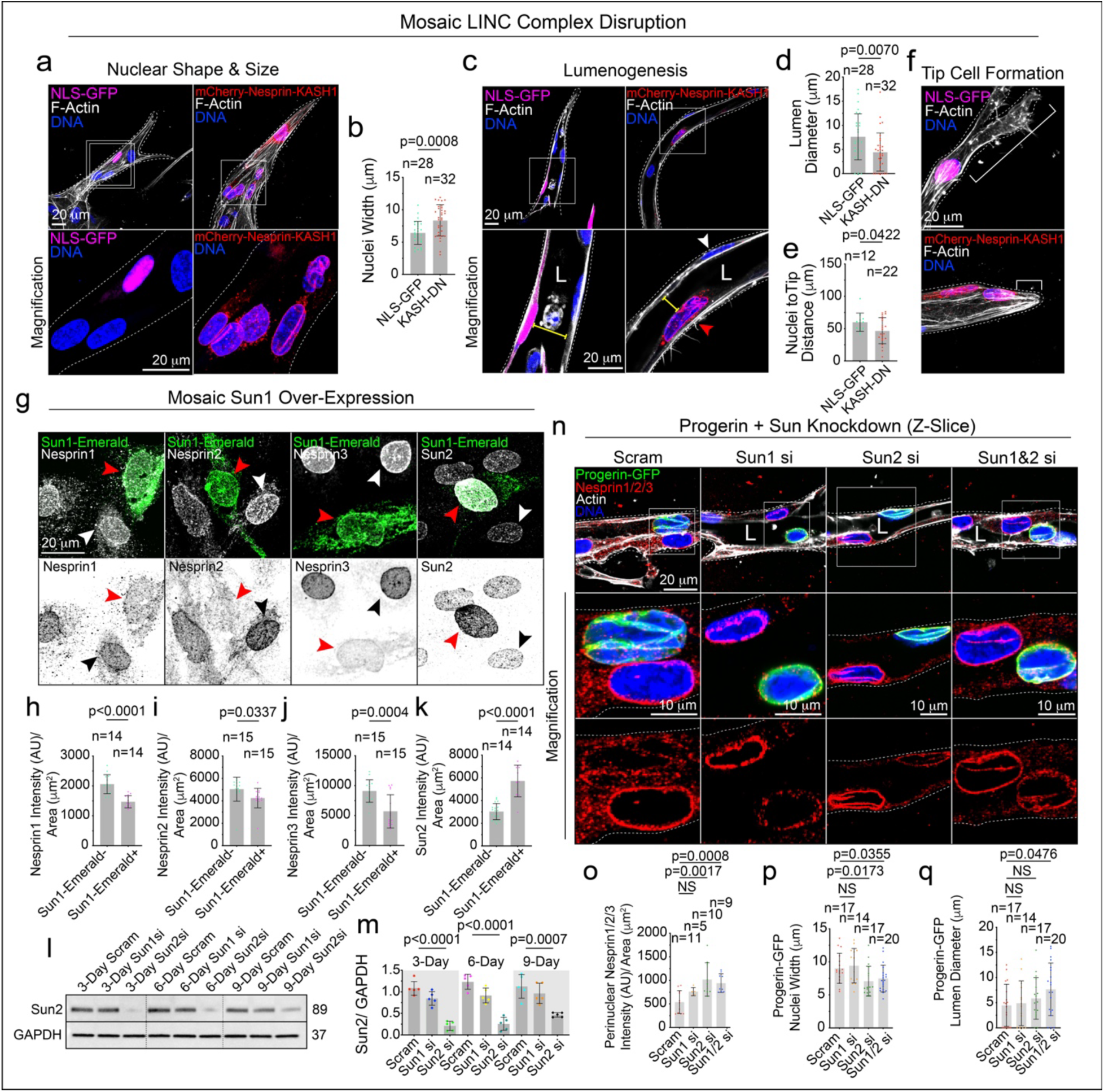
Sun1 Over-Expression Causes Nesprin Displacement and Knockdown Improves Nuclear Shape and Lumen Formation. a) Z-slice image representatives of KASH-domain binding (dominant negative) and control NLS-GFP transfected cells. Quantification of b) nuclear width for control nuclear localized sequence (NLS)-GFP and mCherry-Nesprin-KASH expressing cells. c) Z-slice 60x image representatives of NLS-GFP and mCherry-Nesprin-KASH lumen (L) formation. Quantification of d) lumen diameter at fluorescent cells and e) nuclei to tip distance of fluorescent cells. f) Image representatives of fluorescently positive tip cells. Brackets represent distance quantified g) Image representatives of mosaic Sun1-Emerald expression with Nesprin-1, Nesprin-2, Nesprin-3, and Sun2 staining. Red arrowheads denote Sun1-Emerald positive cell and black arrowhead denotes Sun1-Emerald negative cell. Quantification of fluorescent intensity of h) Nesprin-1, i) Nesprin-2, j) Nesprin-3, and k) Sun2 in Sun1-Emerald negative and positive cells. l) Western blotting for Sun2 knockdown after 3, 6 and 9 days post-electroporation. m) Quantification of western blotting repeats (n=5). n) Image representatives of vessels expressing mosaic Progerin-GFP with universal knockdown of Sun1, Sun2, and double Sun1 and Sun2. Quantification of the Progerin-GFP positive nuclei o) Nesprin1/2/3 intensity p) nuclear width, and q) lumen diameter adjacent the nucleus. For all panels box denotes area of magnification.

Because our previous experiments demonstrated altered SUN protein abundance in Progerin-expressing endothelial cells, and SUN1 has been identified as a major target of Progerin in other cell types,^9^ we next examined how increased SUN1 expression affected LINC complex organization. Mosaic overexpression of a viral construct encoding SUN1-Emerald resulted in a marked reduction of Nesprin-1, Nesprin-2, and Nesprin-3 at the nuclear envelope (**Figure 5g–j**), while increasing SUN2 abundance compared to non-transduced cells (**Figure 5g, k**). These findings suggest that excess SUN may alter LINC complex stoichiometry and contribute to the loss of functional nuclear-cytoskeletal connections observed in Progerin-expressing cells.

We wondered whether correcting excess SUN proteins could restore nuclear localization in Progerin expressing cells. Because complete depletion of SUN proteins abolishes LINC complex assembly, we employed transient siRNA-mediated knockdown followed by an extended recovery period to achieve partial restoration of SUN protein expression in Progerin expressing cells (**Figure 5n-o**). We measured nuclear envelope localization of Nesprin-1, Nesprin-2, and Nesprin-3 in Progerin expressing cells following SUN1, SUN2, or combined SUN1/SUN2 knockdown compared with scramble controls (**Figure 5o**). Restoration of Nesprin localization significantly improved nuclear morphology for SUN2 and SUN1/SUN2 knockdown vessels and increased lumen diameter for Sun1/Sun2 knockdown vessels compared with scramble, Progerin+ vessels (**Figure 5q-s**). In agreement with our results using the KASH construct, these findings indicate that Progerin-induced disruption of the LINC complex contributes to endothelial dysfunction and correcting LINC complex organization can restore normal endothelial behavior.

To better understand how individual LINC complex proteins contribute to endothelial morphogenesis we depleted Lamin-A, SUN proteins, and Nesprins with siRNA. Knockdown of each component individually disrupted endothelial migration, nuclear positioning, and nuclear morphology during vessel formation, although these defects were also consistently less severe than those induced by Progerin expression (**Supplemental Figure 5a–f**). Thus, our data shows that intact LINC complex function is required for endothelial nuclear positioning and lumen formation, and its disruption contributes to nuclear-cytoplasmic disruption in Progeric endothelial cells.

To query the contribution of specific cytoskeletal proteins to nuclear remodeling during endothelial sprouting, we inhibited actin polymerization (Latrunculin A), vimentin organization (FiVe1), or microtubule dynamics (nocodazole) during *in vitro* vessel maturation of HUVECs (**Supplemental Figure 5g**). The number of lumenized sprouts was significantly reduced by treatment with either cytoskeletal inhibitor (**Supplementary Figure 5h**). Microtubule stabilization did not result in aberrantly shaped nuclei. However, nuclei did adopt a 2D planar cell nuclear phenotype; they were flatter and wider compared to all other conditions (**Supplemental Figure 5i**). Vimentin and actin branching inhibition both resulted in misshapen nuclei with significantly reduced sphericity in addition to significantly reduced lumen formation relative to DMSO control vessels (**Supplemental Figure 5i-k**). Western blotting of inhibitor-treated HUVECs confirmed effective pathway inhibition and revealed a significant increase in expression of DNA damage marker H2A.x suggesting possible chromatin damage associated with misshapen nuclei (**Supplemental Figure 5l-o**).^30,37,41^ Furthermore, inhibition of actin polymerization impacted vimentin signal, suggesting an interplay of these proteins. Together, these findings establish that coordinated interactions between the LINC complex and the actin and vimentin cytoskeleton are required for endothelial nuclear remodeling during angiogenic lumen formation.

### Rescue of Progeria Patient Stem Cell Derived Endothelial Defects by Progerinin Treatment

We tested whether restoration of the nuclear-cytoskeletal interface in progeric cells could rescue endothelial sprouting and lumenization. Towards translation to patients, we examined patient-derived endothelial cells generated from induced pluripotent stem cells (iPSCs) derived from a patient with Progeria and the patient’s unaffected parent as a control. These cells were differentiated and cultured in a three-dimensional sprouting assay to assess ex vivo vessel formation (**Figure 6a**). We compared the effects of Lonafarnib, a farnesyltransferase inhibitor and the only FDA-approved treatment for Progeria, with those of Progerinin, a small molecule that selectively binds the C-terminal domain of Progerin and disrupts its interaction with Lamin A while sparing wild-type Lamin-A.^24^ Progerinin extends lifespan in progeric mice and is a promising therapeutic candidate for patients.^24^

**Figure 6.**
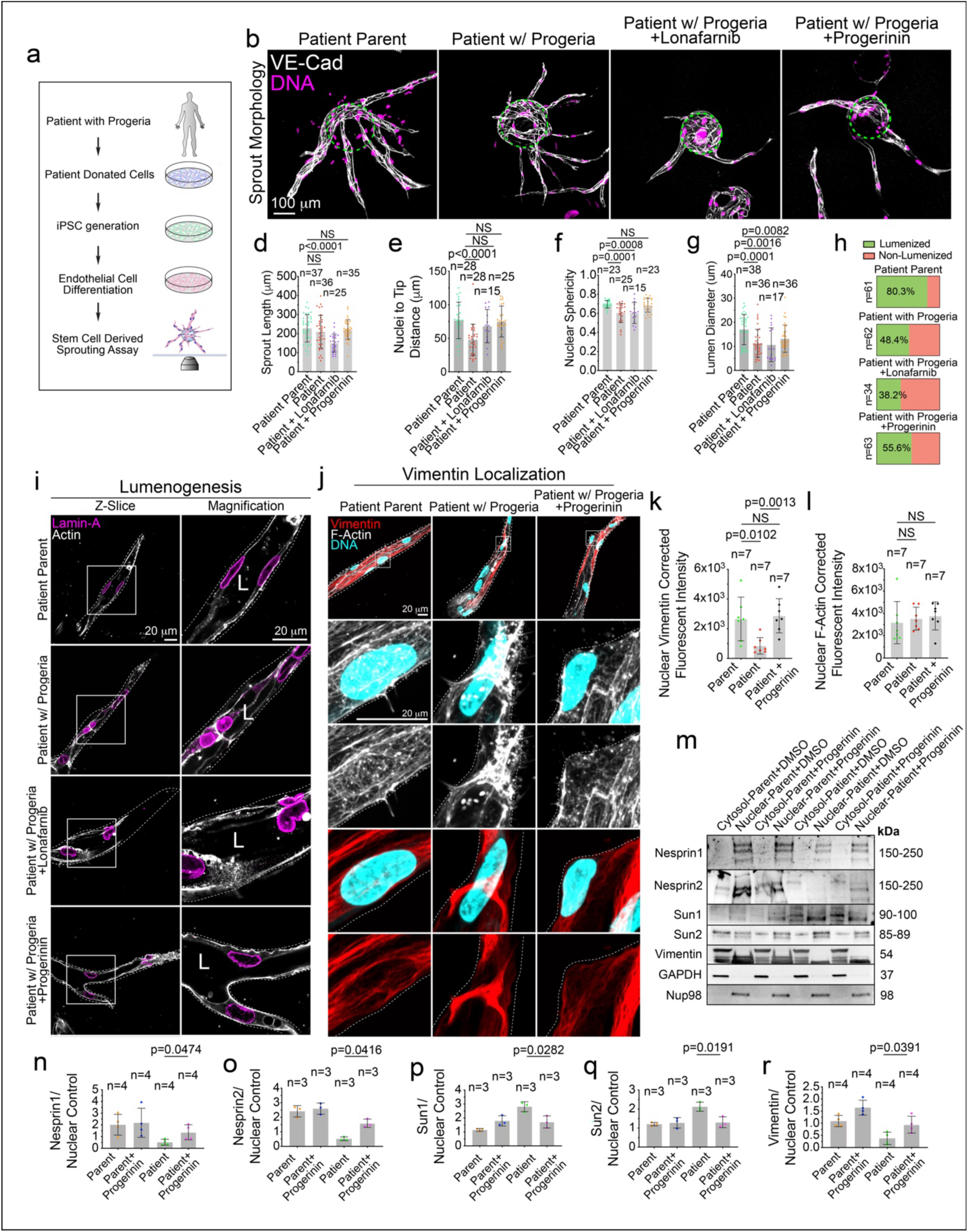
Patient iPSC Derived Vessels are Rescued by Progerinin. a) Cartoon illustrating experimental protocol. b) 20x image representatives of stem cell derived sprouting angiogenesis assay. Dashed line denotes bead. Quantification of c) VE-Cadherin (VE-Cad) positive cells per bead for parent (control) and patient cells, d) sprout length. N=number of vessels, e) nuclei to tip distance, f) nuclear sphericity, g) lumen diameter, and h) percentage of lumenized and non-lumenized vessels. n = number of vessels over 3 separate experiments i) Z-slice image representatives illustrating lumen formation. L denotes lumen. j) Image representatives of F-actin, vimentin, and nuclei. Quantification of nuclear k) vimentin fluorescent intensity and l) f-actin fluorescent intensity. m) Western blotting of cytosolic and nuclear fractions probed for the indicated proteins. n-r) Quantification of western blotting repeats for the indicated proteins.

Analysis of nuclear fractions revealed a significant reduction in nuclear Lamin-A abundance following Lonafarnib treatment, whereas Progerinin did not significantly alter Lamin-A levels compared with vehicle-treated controls (**Supplemental Figure 6a-d**). In contrast, Progerinin treatment reduced nuclear GFP-Progerin abundance in GFP-Progerin-expressing endothelial cells (**Supplemental Figure 6e-f**). These findings are consistent with the distinct mechanisms of action of the two compounds: Lonafarnib inhibits farnesyltransferase, reducing Progerin accumulation at the nuclear envelope by preventing its farnesylation, whereas Progerinin disrupts Progerin-Lamin-A interactions and promotes Progerin degradation while preserving Lamin-A levels..^18,24^

To assess vessel formation, sprouts were stained for VE-cadherin, an endothelial marker, confirming that more than 85% of cells in both parent- and patient-derived cultures were endothelial (**Figure 6b-c**). Sprout length was comparable between parent- and patient-derived vessels, and Lonafarnib treatment significantly impaired sprout growth whereas Progerinin did not (**Figure 6d**). Patient-derived vessels exhibited a significantly shorter distance between endothelial nuclei and the sprout tip than parent-derived vessels, consistent with altered nuclear positioning during sprouting. This defect was improved by both Lonafarnib and Progerinin treatment (**Figure 6e**). Nuclear sphericity differed significantly from controls in untreated and lonafarnib-treated patient cells, but not in Progerinin-treated cells, suggesting that Progerinin treatment improves nuclear morphology. (**Figure 6f**). Relative to parent-derived vessels, lumen formation was significantly impaired in patient-derived vessels and improved following Progerinin treatment (**Figure 6g–i**). Furthermore, perinuclear vimentin localization was markedly reduced in patient-derived endothelial cells and restored following Progerinin treatment (**Figure 6j–k**). We did not assess the effects of Lonafarnib on vimentin organization because protein farnesylation is an important regulator of cytoskeletal dynamics.^40^ Perinuclear f-actin fluorescence intensity did not differ significantly across experimental conditions, although it visually appeared more disorganized in patient vessels compared to parent, and more organized in Progerinin treated versus untreated patient cells (**Figure 6j,l**). Together, these findings demonstrate that patient-derived endothelial cells recapitulate key vascular defects observed in our in vivo and ex vivo models, including altered nuclear positioning, impaired lumen formation, and reduced perinuclear vimentin localization. Several of these defects, particularly those involving nuclear positioning, lumen formation, and vimentin organization, were ameliorated by Progerinin treatment.

Given the improvements in endothelial sprouting and perinuclear vimentin organization following Progerinin treatment, we next investigated whether these effects were accompanied by changes in LINC complex protein abundance. Nuclear fractions were isolated and analyzed for SUN proteins, Nesprins, and vimentin. Nuclear levels of Nesprin-1, Nesprin-2 and vimentin were reduced in patient-derived cells compared to parent cells and increased in patient cells but not parental cells by Progerinin (**Figure 6m-o, r**). Sun1 and Sun2 nuclear expression was increased in patient-derived cells compared to parent, and significantly decreased by Progerinin (**Fig.6m, p, q**). These findings show that Progerinin treatment improved endothelial sprouting and nuclear-cytoskeletal organization, accompanied by changes in LINC complex protein abundance consistent with improved nuclear-cytoskeletal coupling. Collectively, these results support further investigation of strategies targeting the nuclear-cytoskeletal interface as a potential means of improving endothelial function in Progeria and provide a rationale for exploring nuclear-cytoskeletal uncoupling in aging microvessels.

## Discussion

Our findings identify microvascular dysfunction in Progeria and reveal a critical role for nuclear-cytoskeletal coupling in endothelial nuclear positioning and capillary lumen formation. Whereas previous studies of vascular pathology in children with Progeria have largely focused on macrovascular abnormalities and vascular smooth muscle cell loss,^26,36^ our work demonstrates that Progerin also disrupts endothelial function. By altering LINC complex components and the perinuclear cytoskeleton, Progerin impairs endothelial nuclear positioning, flattening, and lumen formation. These findings identify nuclear-cytoskeletal coupling as an important regulator of vascular morphogenesis and suggest that its disruption contributes to microvascular pathology in Progeria. Moreover, the identification of similar endothelial abnormalities in physiologically aged mice suggests that defective nuclear positioning may represent a broader vulnerability of the aging microvasculature.

A central finding of our study is the pronounced loss of perinuclear vimentin organization in Progerin-expressing and aged endothelial cells, particularly during sprouting angiogenesis. This may reflect the extensive cytoskeletal remodeling required for endothelial migration, nuclear repositioning, and lumen formation.^11^ Vimentin intermediate filaments interact with contractile actomyosin arcs that regulate the perinuclear positioning of the vimentin network.^23^ Consistent with this relationship, perturbation of either vimentin or actin altered the localization and phosphorylation state of the other cytoskeletal system in human endothelial vessels, suggesting that their coordinated regulation is important for endothelial nuclear mobility. Notably, aged mouse aortic endothelial cells exhibited similar perinuclear vimentin abnormalities, identifying a shared feature of progeric and aged endothelium. A recent case study also associated a vimentin mutation (1160T>C) with a premature aging phenotype,^10^ further supporting investigation of vimentin dysfunction in age-associated pathology. Although Progeria is distinct from physiological aging, these findings suggest that altered perinuclear cytoskeletal organization may contribute to endothelial dysfunction in both conditions.

Our findings identify disruption of the LINC complex as a potential mechanism underlying the loss of perinuclear cytoskeletal organization in progeric endothelial cells. Previous studies have reported increased SUN1 and SUN2 abundance in progeric cells and explored disruption of SUN-Nesprin interactions as a strategy to mitigate Progerin-associated abnormalities.^6,8,9,17,45^ However, our findings extend these observations by demonstrating reduced Nesprin coverage at the nuclear envelope, altered Nesprin localization, and loss of perinuclear vimentin and vimentin-Nesprin linker, plectin, organization. Together, these alterations suggest that Progerin disrupts the molecular connections linking the nuclear envelope to the cytoskeleton, potentially compromising the cytoskeletal support required for endothelial nuclear remodeling and capillary lumen formation. The mechanisms underlying this disruption remain unclear. Increased cytoplasmic Nesprin signal (Figures 6f and 7h) is consistent with the endoplasmic reticulum-associated Nesprin in fibroblasts reported by Chen et al. in Progerin-expressing fibroblasts.^9^ Importantly, our perturbation experiments demonstrate that depletion of individual LINC complex components also results in nuclear abnormalities and impaired vessel formation, suggesting that simply disrupting nuclear–cytoskeletal connections is insufficient to preserve endothelial function and that maintaining appropriate LINC complex organization is critical for nuclear remodeling.

Importantly, our findings extend beyond Progeria, identifying alterations in nuclear-cytoskeletal components during physiological aging. Aged mouse aortic endothelium exhibited overlapping abnormalities in LINC complex components, while analysis of human endothelial aging revealed distinct transcriptional changes in nuclear structural proteins and their cytoskeletal linkers. In cardiac endothelial cells, increased Lamin-A and vimentin expression occurred alongside reduced expression of Nesprin-encoding genes, suggesting that the transcriptional regulation of nuclear structural components and their connections to the cytoskeleton may become discordant with age. Furthermore, enrichment of nuclear envelope organization-related gene sets was particularly evident in aged cardiac endothelium, highlighting tissue-dependent differences in nuclear architecture-associated transcriptional responses to aging. These findings suggest that the nuclear-cytoskeletal interface represents a potential site of vulnerability in both Progeria and physiological aging, although the molecular changes underlying this vulnerability may differ across conditions and vascular beds.

Experimental disruption of LINC complex function only partially reproduced the endothelial abnormalities induced by Progerin, supporting a functional contribution of impaired nuclear-cytoskeletal coupling to defective vascular morphogenesis. However, the less severe phenotype suggests that additional mechanisms may contribute. One possibility is increased nuclear stiffness, previously reported in fibroblasts derived from patients with Progeria.^43^ Because endothelial nuclei must repeatedly deform and reposition during migration and lumen formation,^7^ increased stiffness could further restrict nuclear movement. These mechanisms are mechanically distinct: LINC complex disruption compromises cytoskeletal force transmission, whereas nuclear stiffening reduces nuclear deformability, which is influenced by Lamin A/C organization and phosphorylation-dependent remodeling.^31,43^ Future studies should determine how these mechanisms interact to regulate endothelial nuclear positioning and whether altered nuclear mechanics contributes to microvascular dysfunction during physiological aging.

Our findings also have potential therapeutic implications for Progeria. The abnormalities observed in developing progeric vasculature suggest that Progerin interferes with vascular morphogenesis early in life, highlighting the importance of investigating interventions that preserve endothelial function. Lonafarnib, the only FDA-approved therapy for Progeria, inhibits farnesylation of multiple cellular proteins and is not specific to Progerin.^32^ In contrast, Progerinin is a small-molecule inhibitor designed to disrupt the interaction between Progerin and Lamin-A.^24,39^ Previous studies demonstrated beneficial effects in progeric mouse models,^24^ and early clinical studies have evaluated its safety and tolerability, although clinical efficacy has not been established.^38^ In our patient-derived endothelial models, Progerinin improved cell expansion, sprouting, and lumen formation, whereas Lonafarnib impaired sprouting under the conditions examined. These findings demonstrate that aspects of Progerin-associated endothelial dysfunction are pharmacologically modifiable and support further investigation of therapeutic strategies targeting nuclear dysfunction.

Collectively, our study identifies endothelial nuclear-cytoskeletal dysfunction as an important contributor to microvascular pathology in Progeria and establishes nuclear positioning and morphology as critical determinants of capillary lumen formation. The identification of related abnormalities in aged endothelial cells suggests that disruption of perinuclear cytoskeletal organization may also contribute to vascular dysfunction during physiological aging, although the underlying mechanisms may differ across tissues and disease contexts. Future studies investigating how LINC complex composition, perinuclear vimentin organization, and nuclear mechanics interact to regulate endothelial function may reveal new strategies to preserve microvascular integrity in Progeria and aging.

## Methods

### Reagents

All reagents, drugs, siRNA, and plasmid information are listed tables 1-4 of associated supplementary information.

### Cell Culture

HEK cells were purchased from ATCC and cultured in Dulbecco’s Modified Medium (DMEM) supplemented with 10% fetal bovine serum. Normal Human Lung Fibroblasts (NHLF’s) were purchased from Lonza and cultured in Lonza FGM-2 Fibroblast Growth Medium-2 Bullet Kit^®^. Pooled Human umbilical vein endothelial cells (HUVECs) were purchased from PromoCell and cultured in proprietary media for 2–5 passages. Isolated mouse lung endothelial cells, as well as iPSC-derived endothelial cells, were cultured in PromoCell EGM-2 for 1-2 passages. All cells were maintained in a humidified incubator at 37 °C and 5% CO2. All medias contained 1% penicillin/streptomycin and were used within 2-3 weeks of opening.

Small interfering RNA was resuspended to a 20 µM stock concentration and used at 0.5 µM. SiRNA was introduced into primary HUVEC’s using electroporation with the Neon^®^ transfection system (ThermoFisher). *In vitro* drug treatment concentrations are provided in Supplementary Table 3.

### iPSC Differentiation

Patient iPSC’s were acquired from the Progeria Research Foundation cell and tissue bank. We utilized cells belonging to the same donors (HGADFN167 iPS 1J, HGFDNFN168 iPS1 D2) for all experimental replicates. Differentiation of iPSC’s was achieved following the protocol described in the Stem Cell Technologies STEMdiff™ Endothelial Differentiation Kit. Following differentiation, cells were split once prior to experimentation and not passaged more than two times in total. Experimental replicates for all iPSC-derived endothelium are the result of 3 separate differentiations from 3 vials received.

### Mouse Endothelial Cell Isolation

Endothelial cell isolations were achieved by harvesting the lung of adult mice at 20-weeks and 104-weeks (aged) of age. Following euthanasia, lung tissue was quickly removed and transported to a sterile hood on a tissue culture dish. The lung was mechanically reduced using a razorblade, followed by resuspension with ice cold DMEM and pipetteing with progressively smaller syringes to continue to break up lung tissue. Following mechanical reduction, homogenate was pelleted and resuspended with type II collagenase/dispase and digested for 1 hour with DNAse. The reaction was stopped with ice-cold DMEM, and digestion was passed through a cell strainer.

After digestion, lysate incubated in CD45 coated beads (for removal of immune cells which are CD31 positive). After CD45 bead removal with magnetic separator, lysate was incubated with CD31 coated beads to isolate the endothelium. Upon isolation cells were incubated in EGM-2 media supplemented with puromycin (10ug/mL). Media was changed on isolations 4 hours after plating.

### In Vitro & Ex Vivo Sprouting Angiogenesis Assay

Sprouting angiogenesis/fibrinogen-bead assays were performed as reported by Nakatsu et al.^38^ HUVECs were coated onto Cytodex-3 microcarrier beads (Cytvia) over 4 hours and plated overnight. SiRNA-treatment or viral transduction was performed the same day the beads were coated (Supplementary Table 1). Coated microbeads were embedded in a fibrinogen/aprotinin matrix in a 10cm live imaging dish. Once the clot was formed media was overlaid along with 100,000 NHLFs. Media was changed every other day along with monitoring of sprout development. Fixation occurred after 7 days post embedding. *Ex Vivo* and iPSC-derived sprouting assay protocols did not differ; however, both assays were extended from 7 days to 10 days.

Mosaic siRNA sprouting assays were achieved by treating a population of cells with Cell Tracker (ThermoFisher) and immediately electroporating siRNA into only cell tracker treated cells. These cells are then combined with untreated cells 50:50 and promptly coated onto beads the same day.

Sprout numbers were determined by counting the number of multicellular sprouts emanating from an individual microcarrier beads across multiple beads in each experiment. Sprout lengths were determined by measuring the length of a multicellular sprout beginning from the tip of the sprout to the microcarrier bead surface across multiple beads. Percent of lumenized sprouts were determined by quantifying the proportion of multicellular sprouts whose length (microcarrier bead surface to sprout tip) was less than 70% lumenized across multiple beads. Lumen diameters were determined by two methods, 1) in mosaic experiments (cell tracker or mosaic transductions) lumen diameter was measured at the location of the nucleus of the positive cell within the vessel stalk and 2) in experiemnts in which all cells were altered (siRNA knockdowns) diameters were measured at the widest point/centrally in the vessel sprout. Experimental repeats are defined as an independent experiment in which multiple cultures, containing numerous sprouting beads were quantified; this process of quantifying multiple parameters across many beads and several cultures was replicated on different days for each experimental repeat.

### Lentivirus Generation & Transduction

Lentiviral plasmids were either purchased from Addgene or cloned by using the LR Gateway Cloning method. For lentiviral generation genes of interest and fluorescent proteins were isolated and incorporated into a pME backbone via Gibson reaction. Following confirmation of the plasmid by sequencing the pME entry plasmid was mixed with the destination vector and LR Clonase. The destination vector used in this study was pLEX 307. Once validated, the destination plasmids were transfected with the three required viral protein plasmids: pVSVG (gift from Bob Weinberg; Addgene plasmid #8454) and psPAX2 (gift from Didier Trono; Addgene plasmid #12260) into HEK 293 cells. The transfected HEKs had media changed 4 h post transfection. Transfected cells incubated for 3 days, and virus was harvested by virus concentration media and a 1-hour centrifugation in a fixed rotor at 4 Celsius.^22^

### Cell Stimulation Assay

Treated HUVEC’s were lysed at baseline, serum starvation, or 10 minutes stimulated. For baseline, cells were lysed without intervention, maintained in EGM-2 media. Serum-starvation, and stimulation, consisted of 2 hours incubation with OPTI-MEM. Following starvation cells were either lysed (starved condition) or received a media change with EGM-2 and allowed to incubate for 10 minutes (stimulated condition). Cells were lysed as described in immunoblotting protocol.

### Mouse Husbandry

Mice for this investigation were housed in the Yale Animal Resources Center (YARC) under a 12-hour light/dark cycle at an ambient temperature of 20-24 °C. The mouse strain used throughout the study was C57BL/6-Tg(LMNA*G608G) and was purchased from Jackson Laboratories (Strain #: 010667). The donating investigator to the Jackson Laboratory was Dr. Francis Collins of the National Institute of Health. Offspring were genotyped by standard PCR from tail or ear DNA. All experimental procedures were reviewed and approved by IACUC.

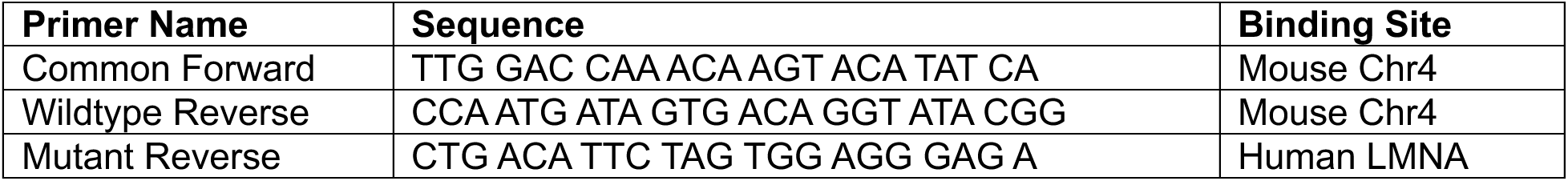

### Developmental Angiogenesis & Vascular Profiling

Neonatal pups were euthanized by decapitation at P6 for developmental angiogenesis analysis of mouse retinas. P6 eyes were fixed with 4% paraformaldehyde (PFA) and constant agitation for 10 minutes, washed and promptly dissected for immunohistochemistry (IHC). Profiling of the adult vasculature was achieved by euthanizing 20-week, or 10-week, mice with isoflurane. Adult retinas were fixed with 4% PFA for 1 hour with constant agitation and retinas were dissected promptly following fixation.

Vascular perfusion assays for pup and adult mice were achieved by intraperitoneal injection of Sulfo-NHS-Biotin (5mg/g of body weight) resuspended in PBS. Following resuspension, Sulfo-NHS-Biotin was used the same day and not stored for later use. Incubation required 1-hour (pups) or 2-hours (adults) before euthanasia. Sulfo-NHS-Biotin was detected with IHC by Alexa Fluor-conjugated streptavidin.

Male and female mice were used for all experiments, and no significant sex differences were evident in our quantification.

### Live and Fixed Microscopy

All fluorescence microscopy was performed on an inverted Nikon Eclipse Ti2 microscope (Nikon Instruments) with a Perfect Focus System, equipped with a Dragonfly unit. Lasers were coupled into a multimode fiber going through the Andor Borealis unit reshaping the beam from a Gaussian profile to a homogenous flat top. As dichroic mirror, a CR-DFLY-DMQD-01 was used. Fluorescence light was spectrally filtered with an emission filter (TR-DFLY-F600-050) and imaged with a scientific complementary metal oxide semiconductor (sCMOS) camera (Sona 4BV6X, Andor Technologies). For live imaging, the microscope was equipped with a Okolab live imaging chamber and an internal environment of 5% CO2 and 37 °C was maintained.

Image processing was performed on FIJI (Image J) and IMARIS. Nuclei were segmented and 3D modeled using the IMARIS Surface Rendering feature. Live imaging analysis of nuclear movement was achieved by modeling nuclei with the surface rendering tool and running advanced statistics in IMARIS which provided sphericity, track straightness, track length, displacement and volume change data.

### In Vitro & In Vivo Immunostaining

For immunofluorescence imaging, 2-dimensional HUVEC assays were fixed with 4% PFA for 7 min. ECs were then washed three times with PBS and permeabilized with 0.5% Triton-X (Sigma) for 10 min. After permeabilization, cells were washed three times with PBS. ECs were then blocked with 2% bovine serum albumin (BSA) for 30 min. Once blocked, primary antibodies were incubated for 2 hours, washed thrice and incubated with secondary antibodies for 1 hour. Washed and imaged. 3-dimensional sprouting assays were fixed for 40 minutes, washed three times and permeabilized for 1-2 hours (antibody dependent). Following permeabilization, sprouts were blocked for a minimum 1-hour before overnight primary antibody incubation. Thereafter, primary antibodies were removed, and the cells were washed 3 times with PBS. Secondary antibody with 2% BSA were added and incubated overnight, washed 3 times with PBS, and imaged. All primary and secondary antibodies are listed in Supplementary Table 3.

Following fixation, pup and adult retinas were washed and blocked with *in vivo* blocking buffer (1% Triton-X100, 5% BSA, and 5% donkey serum in PBS) for 1 hour. After blocking, retinas were incubated in primary antibody for 4 hours, or overnight, washed 3-5 times (antibody dependent) and treated with secondary for 4 hours. Retinas were then washed and mounted for imaging. Fluorescent imaging of proliferative cells was achieved with the Click-iT™ EdU Cell Proliferation Kit (Invitrogen) and provided protocol.

Adult aortas were isolated and trimmed of surrounding adipose. Aortas were fixed with 4% PFA for 20 minutes and blocked with *in vivo* blocking buffer for 30 minutes. Primary antibody incubated for 4 hours, aortas were washed 3 times, then secondary antibody treatment for 2-3 hours (antibody dependent) followed by washes and mounting.

### In Vitro & In Vivo Immunoblotting

HUVEC’s from stimulation assays and knockdown confirmations were lysed with cell scraper using RIPA buffer containing 1× PhosSTOP™ tablets (Sigma) and cOmplete™, Mini, EDTA-free Protease Inhibitor Cocktail tablets (Sigma). Cell lysate was vortexed and centrifuged for 10 minutes and supernatant was transferred to new tube.

Western blotting of HUVEC and iPSC-derived endothelial cell nuclear and cytosolic fractions were achieved by NE-PER™ Nuclear and Cytoplasmic Extraction Reagents kit.

Mouse retina lysate was acquired by removal and timely dissection of the retina in PBS. Retinas were gently transferred to a microcentrifuge tube and excess PBS removed. Both retinas from a single mouse were combined and lysed with the previously described RIPA cocktail. Lysate was vortexed, centrifuged for 10 minutes, and supernatant transferred to a new vial.

For all specimens, lysate was brought up with Lamelli and incubated in a heat block at 98 degrees for 3-4 minutes. All samples were used within one week of preparation.

### scRNA-Seq Analysis of Endothelial Aging

To investigate age-associated transcriptional changes in nuclear architecture-related genes, human endothelial single-cell and single-nucleus RNA-sequencing data were obtained from the Human Aging Endothelial Cell Atlas (HAECA, GSE305900).^14^ The atlas comprises 375,829 endothelial cells from 290 samples across 12 human tissues: adipose tissue, brain, breast, heart, intestine, kidney, liver, lung, skeletal muscle, ovary, skin, and uterus.

To account for donor-level biological variability and avoid pseudo replication, RNA counts were aggregated by donor within each tissue, generating pseudo-bulk expression profiles. Pseudo-bulks containing fewer than 20 endothelial cells were excluded, and differential expression analyses required at least three independent donors per age group. The primary analysis was performed separately for each tissue using the limma–voom framework. Lowly expressed genes were filtered, library sizes were normalized using the trimmed mean of M-values (TMM) method. Differential expression between aged and young donors was estimated using linear models with empirical Bayes moderation.

To assess coordinated transcriptional changes in nuclear architecture-related pathways, pre-ranked gene set enrichment analysis (GSEA) was performed independently for each tissue using moderated t statistics from the limma–voom analysis. Gene Ontology (GO) Biological Process and Cellular Component gene sets were obtained from MSigDB and enrichment was calculated with fgsea. Pathways related to nuclear envelope organization, nuclear lamina, nuclear membrane, nuclear pore complexes, and intermediate filament organization were selected for visualization.

For the EC analysis, 22 donors (10 young and 12 aged) were included. Genes annotated to the GO cellular component term nucleus (GO:0005634), including descendant terms, were identified using org. Hs.eg.db and AnnotationDbi and intersected with the cardiac differential expression results. The resulting 5,328 nucleus-annotated genes were ranked according to their limma–voom log_2_FC (aged versus young). Statistical significance from gene-wise p values were adjusted using the Benjamini–Hochberg (BH) procedure across the set of genes tested in cardiac ECs.

To investigate tissue-specific heterogeneity in the age-associated expression of genes involved in nuclear lamina organization (LMNA, LMNB1), LINC complex function (SUN1, SUN2, SYNE1– 4), and cytoskeletal coupling (VIM, PLEC), we performed an independent differential expression approach using donor-level pseudo-bulk counts using the edgeR quasi-likelihood framework. Estimated log_2_FC for the 10 predefined nuclear lamina–LINC–cytoskeleton genes were visualized across the 12 tissues in a hierarchically clustered heatmap. Significance was based on BH-adjusted P values across all genes tested within each tissue.

### Statistical Analysis

*In vitro* experiments were repeated a minimum of three times*. In vivo* biological replicates were a minimum of 3, but mostly 5 or more. Statistical analysis and graphing were performed using GraphPad Prism. Statistical significance was assessed with a student’s unpaired t-test for a two-group comparison. Multiple group comparisons were carried out using a one-way analysis of variance (ANOVA) followed by a Dunnett multiple comparisons test. Data was scrutinized for normality using Kolmogorov–Smirnov test. Statistical significance set a priori at p < 0.05.

### Ethical Consideration

All research presented followed guidelines outlined by the Yale University Institutional Biosafety Committee (IBC) and Institutional Animal Care and Use Committee (IACUC).

## Notes

### Competing Interest Statement

The authors have declared no competing interest.

### Summary of Updates

All figures and text have been updated.

