## Supplementary Materials for "Nuclear–Cytoskeletal Uncoupling in Aging Capillaries Disrupts Endothelial Nuclear Flattening and Lumen Formation"

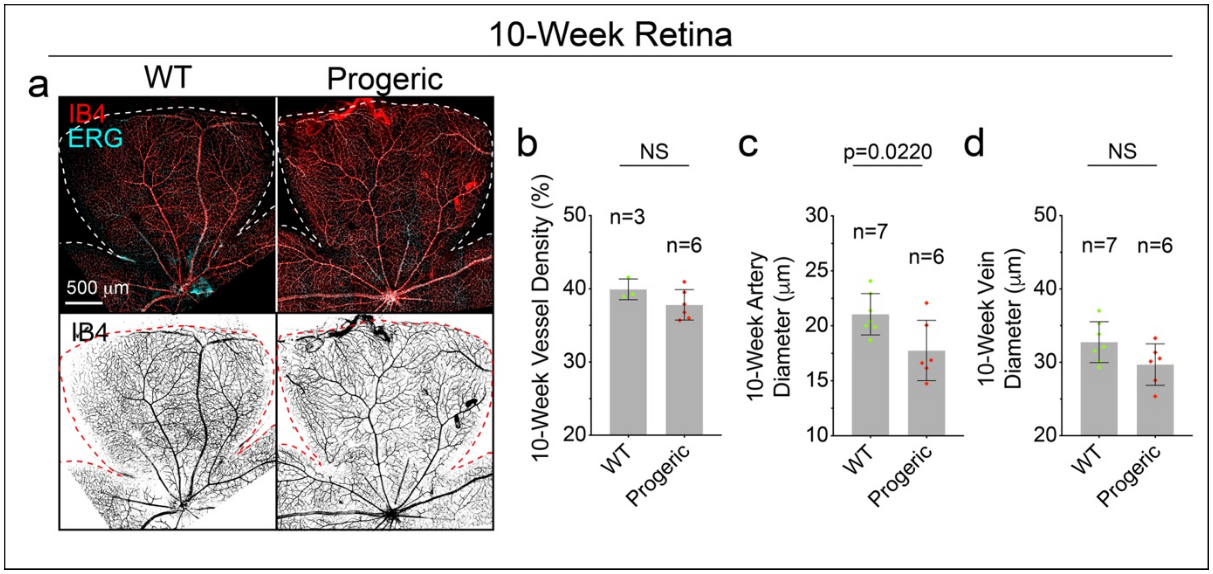


**Supplemental Figure 1. Progerin Expression Delays Development in P6 Pups.** a) Image representatives of 10-week wildtype and progeric retina vasculature marked by IB4 and endothelial nuclear marker, ERG. Quantification of 10-week b) vessel density, c) artery diameter, and d) vein diameter. NS= Not Significant.

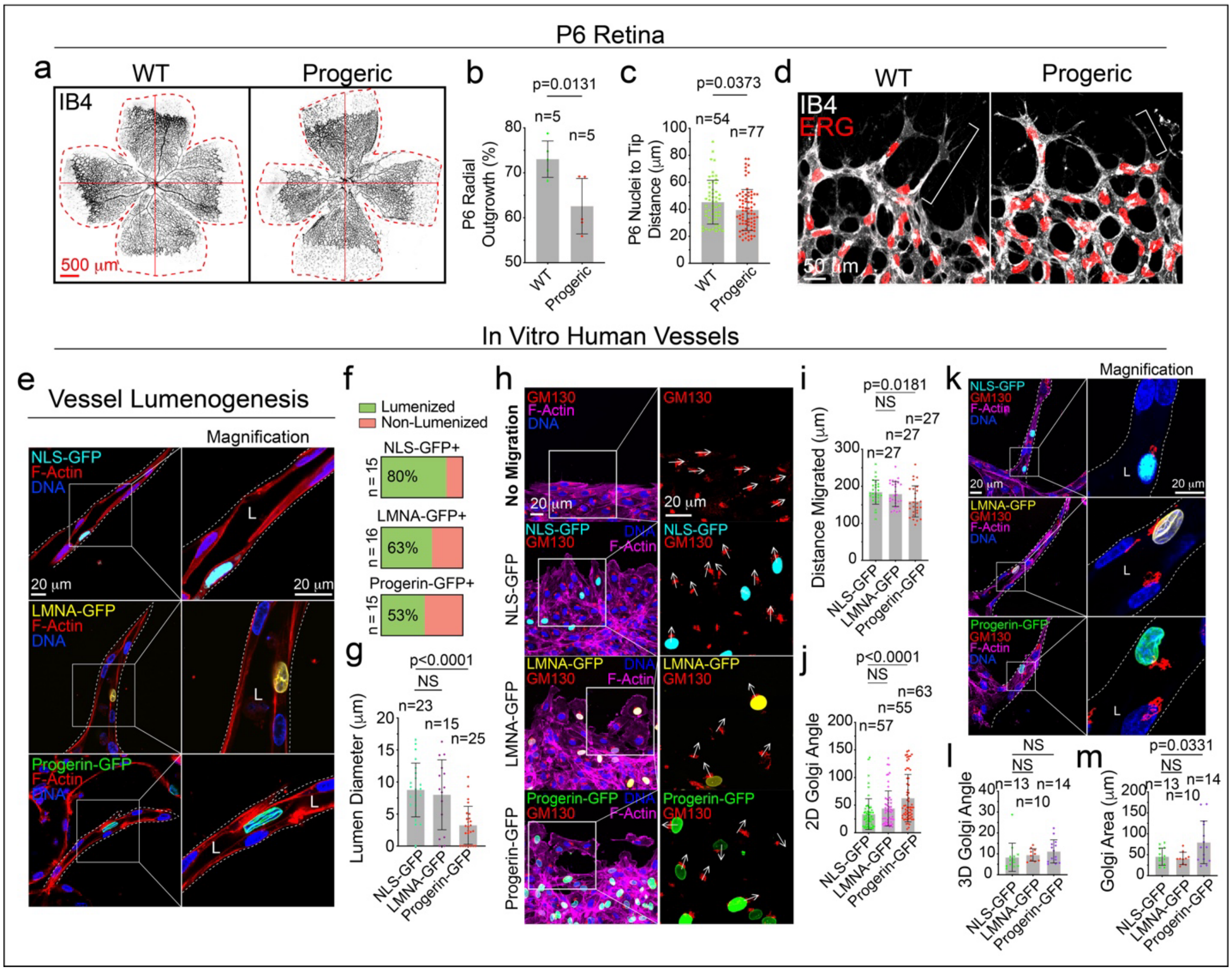


**Supplemental Figure 2. Altered Polarity in Progeric Endothelial Cells.** a) Image representatives of P6 retina vascular outgrowth marked by IB4. Dashed line denotes retina outline. Quantification of b) P6 retina outgrowth and c) P6 nuclei to tip distance. d) Image representatives of vascular front with vascular marker, IB4, and endothelial nuclear marker, ERG. Brackets illustrate cellular projection length. e) 60x Z-slice image representatives for indicated conditions. Quantification of f) percent lumenization and g) lumen diameter measured at GFP-positive cell nuclei within the vessel shaft. h) Image representatives of pre-migration (top row) and post-5 hours of migration for the indicated conditions. Arrows indicate directionality of Golgi marked by GM130. Quantification of i) distance migrated and j) Golgi angle relative to the direction of migration. k) Image representatives of Golgi orientation in vessel sprouting assay for indicated conditions. Quantification of l) Golgi angle in vessel sprouts relative to the tip cell and m) Golgi area. For all panels box denotes area of magnification, L denotes lumen, and NS = not significant.

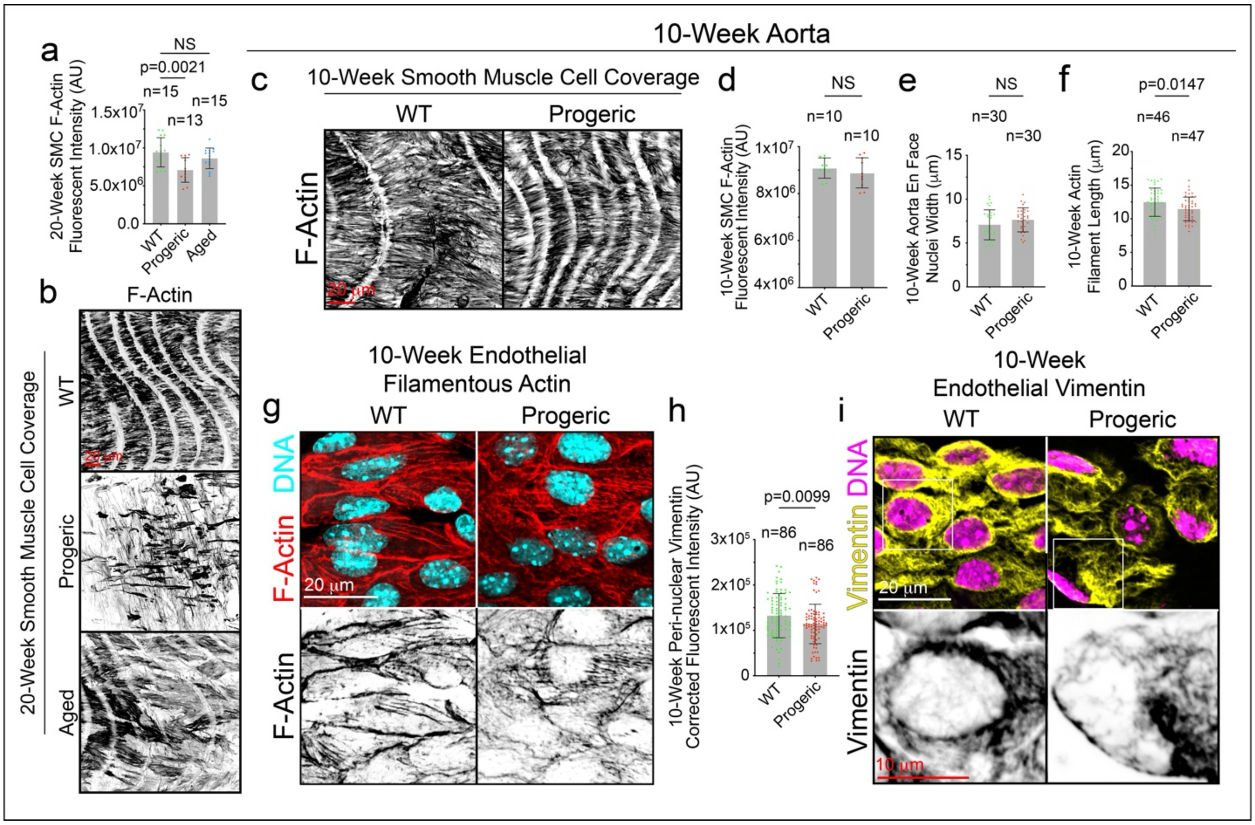


**Supplemental Figure 3. Aorta Endothelium Smooth Muscle Cell and Cytoskeletal Defects**
a) Quantification of smooth muscle cell coverage by intensity of filamentous actin in 20-week mice. b) 60x image representatives of 20-week-old aortic smooth muscle cell coverage marked by f-actin. c) 60x image representatives of 10-week-old aortic smooth muscle cell coverage marked by f-actin. Quantification of d) f-actin intensity, e) nuclear width, f) endothelial actin filament length in the aorta. g) Image representatives of 10-week aorta endothelial f-actin. h) Quantification of peri-nuclear vimentin intensity. i) Image representatives of 10-week aortic endothelial vimentin. Box denotes area of magnification. NS = Not Significant.

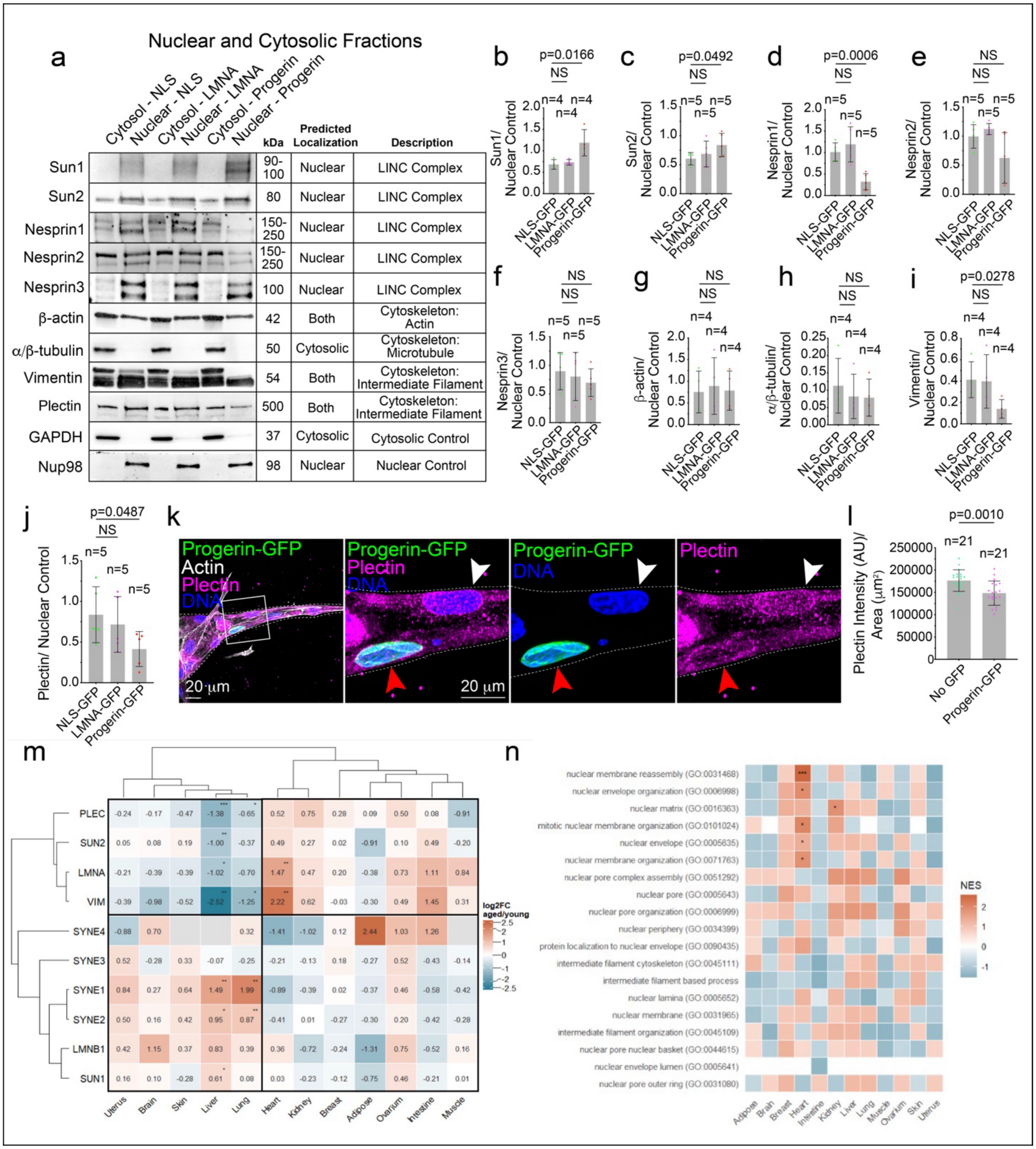


**Supplemental Figure 4. Expression of LINC complex proteins in progeric endothelium and RNA levels in aged human endothelium.** a) Western blotting of nuclear and cytoplasmic fractions for described LINC complex proteins and cytoskeletal proteins. The first fraction is nuclear localized GFP expressing cells, second is full length Lamin-A expressing cells and last is Progerin expressing cells. Nesprin-1 and -2 probes resulted in bands ranging from 150kDa beyond the scope of our protein ladder, bands shown here are within the limit of the protein ladder. b-j) Quantification of western blotting repeats. Nuclear control refers to Nup98. Nesprin-1 and -2 probes resulted in multiple bands; all bands were quantified. NS = Not Significant. k)

Image representatives of vessels mosaically expressing Progerin-GFP stained for Plectin. White arrowhead denotes Progerin-negative cell and red arrowhead denotes Progerin-positive cell. l) Quantification of perinuclear Plectin. m) Clustered heatmap showing age-associated differential expression of 10 genes involved in nuclear lamina organization (*LMNA, LMNB1*), LINC complex components (*SUN1, SUN2, SYNE1, SYNE2, SYNE3, SYNE4*), and cytoskeletal coupling (VIM, PLEC) across human tissues. Differential expression was assessed using donor-level endothelial pseudobulk analysis with the edgeR quasi-likelihood. Cell colors and numerical labels represent increased (red), decreased (blue) or unavailable (gray) $\log_2$FC (aged versus young). Genes and tissues were hierarchically clustered using Pearson correlation distance and average linkage. BH-adjusted significance across genes tested within each tissue (*FDR < 0.05, **FDR < 0.01, ***FDR < 0.001). Gray cells indicate unavailable differential expression estimates. n) Heatmap showing GSEA of GO terms associated with nuclear envelope organization, nuclear lamina, nuclear membrane, nuclear pore complexes, and intermediate filament organization across ECs from different human tissues. Colors represent positive (red) or negative (blue) normalized enrichment scores (NES) in aged relative to young endothelial cells. * Benjamini–Hochberg (BH) corrected FDR < 0.05, **FDR < 0.01, ***FDR < 0.001.

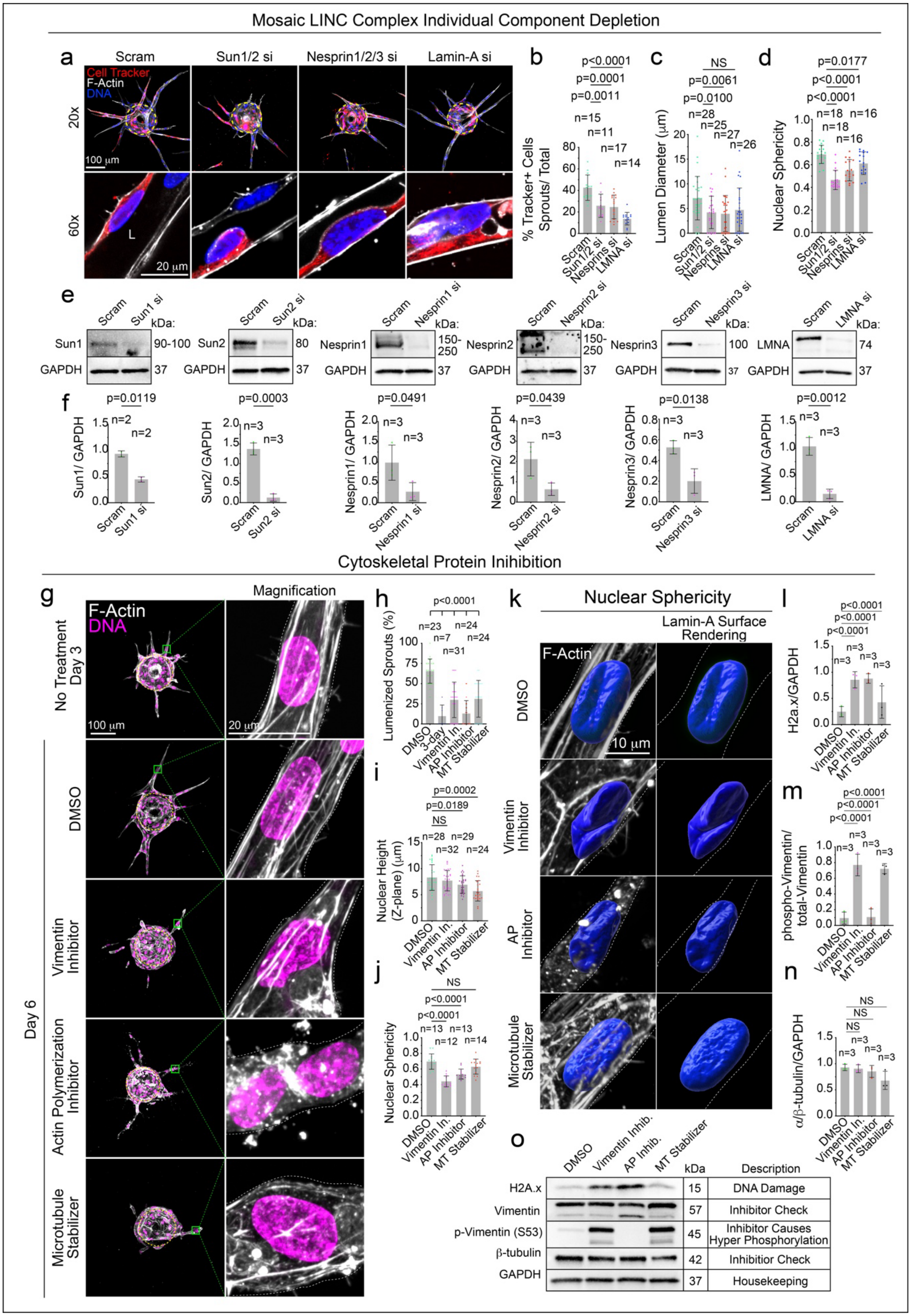
Mosaic LINC Complex Individual Component Depletion
a
Scram
Sun1/2 si
Nesprin1/2/3 si
Lamin-A si
Cell Tracker
F-Actin
DNA
20x
60x
100 μm
20 μm
L
b
p<0.0001
p=0.0001
p=0.0011
n=15
n=11
n=17
n=14
% Tracker+ Cells Sprouts/ Total
Scram
Sun1/2 si
Nesprins si
LMNA si
c
NS
p=0.0061
p=0.0100
n=28
n=25
n=27
n=26
Lumen Diameter (μm)
d
p=0.0177
p<0.0001
p<0.0001
n=18
n=16
n=18
n=16
Nuclear Sphericity
e
Scram
Sun1 si
kDa:
Sun1
90-100
GAPDH
37
Sun2 si
Sun2
80
Nesprin1 si
Nesprin1
150-250
Nesprin2 si
Nesprin2
Nesprin3 si
Nesprin3
100
LMNA si
LMNA
74
f
p=0.0119
n=2
Sun1/ GAPDH
p=0.0003
n=3
Sun2/ GAPDH
p=0.0491
Nesprin1/ GAPDH
p=0.0439
Nesprin2/ GAPDH
p=0.0138
Nesprin3/ GAPDH
p=0.0012
LMNA/ GAPDH
Cytoskeletal Protein Inihibition
g
Magnification
F-Actin
DNA
No Treatment Day 3
100 μm
20 μm
Day 6
DMSO
Vimentin Inhibitor
Actin Polymerization Inhibitor
Microtubule Stabilizer
h
p<0.0001
n=23
n=7
n=31
n=24
n=24
Lumenized Sprouts (%)
DMSO
3-day
Vimentin In.
AP Inhibitor
MT Stabilizer
i
p=0.0002
p=0.0189
NS
n=28
n=29
n=32
n=24
Nuclear Height (Z-plane) (μm)
j
NS
p<0.0001
p<0.0001
n=13
n=12
n=13
n=14
Nuclear Sphericity
k
Nuclear Sphericity
Lamin-A Surface Rendering
F-Actin
10 μm
DMSO
Vimentin Inhibitor
AP Inhibitor
Microtubule Stabilizer
l
p<0.0001
p<0.0001
p<0.0001
n=3
H2a.x/GAPDH
m
p<0.0001
p<0.0001
p<0.0001
phospho-Vimentin/ total-Vimentin
n
NS
NS
NS
α/β-tubulin/GAPDH
o
DMSO
Vimentin Inhib.
AP Inhib.
MT Stabilizer
kDa
Description
H2A.x
15
DNA Damage
Vimentin
57
Inhibitor Check
p-Vimentin (S53)
45
Inhibitor Causes Hyper Phosphorylation
β-tubulin
42
Inhibitior Check
GAPDH
37
Housekeeping

**Supplemental Figure 5. Loss of LINC Complex Proteins Cause Nuclear Obstruction and Lumen Collapse.** a) 20x and 60x image representatives of described mosaic siRNA knockdown (KD) cells. Quantification of b) sprouts per bead, c) sprout length, and d) Percent cell tracker-positive cells in sprouts by total cell tracker-positive cells on the bead and sprouts. e) Western blotting for knockdown confirmation of described proteins. f) Quantification of western blotting replicates. g) Image representatives of vessels before and after inhibitor treatment. DMSO = vehicle, Vimentin inhibitor = Five1, actin polymerization inhibitor = Latrunculin A, and Microtubule stabilizer = nocodazole. Column one is 20x vessel morphology, column two is z-projection of 60x morphology, and third column is magnification. Box denotes area of magnification. Quantification of h) nuclear length relative to vessel length, i) nuclear width and j) percent lumenized vessels. k) Western blotting for indicated proteins. l-n) Quantification of western blotting repeats for indicated proteins.

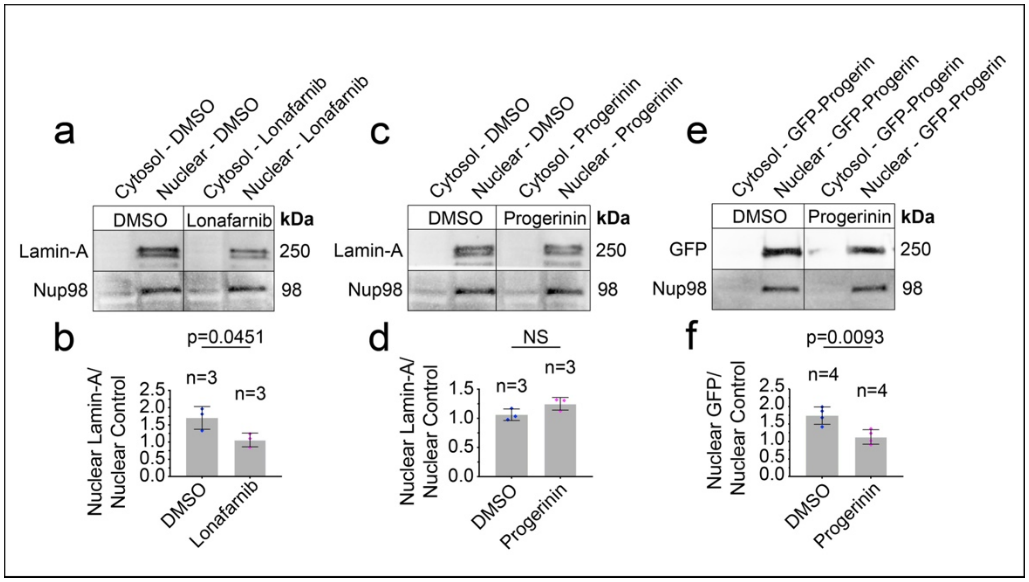


**Supplemental Figure 6. Progerinin is Progerin Specific Inhibitor.** Western blotting of cytosolic and nuclear fractions for a) DMSO and Lonafarnib treated HUVECs probed for Lamin-A and nuclear control Nup98, b) DMSO and Progerinin treated HUVECs probed for Lamin-A and Nup98, and c) DMSO and Progerinin treated HUVECs transduced with Progerin-GFP and probed for GFP and Nup98. Quantification of d) blotting replicates from figure a, e) blotting replicates from figure b, and f) blotting replicated from figure c. NS = Not Significant.

**Supplementary Table 1. Major Resource Table.**

| Reagent | Vendor | Catalog # |
|---|---|---|
| OPTI-MEM 1 Reduced Serum Medium, no phenol red | Thermo Scientific | 31985070 |
| Polyethyleneamine Branched (PEI) Table 1. | Sigma-Aldrich | 408727 |
| Chloroquine Diphosphate Crystalline (CQ) | Sigma-Aldrich | C6628-25G |
| Normal Human Lung Fibroblasts (NHLF's) | Lonza | CC-2512 |
| Human Embryonic Kidney Cells (HEK 293-A) | Thermo Scientific | R70507 |
| Human Umbilical Veinous Endothelial Cells (HUVEC's) | PromoCell | C-12203 |
| Endothelial Cell Growth Medium 2 | PromoCell | C-22011 |
| FGM™-2 Fibroblast Growth Medium-2 BulletKit™ | Lonza | CC-3132 |
| DMEM, High Glucose, with L-Glutamine | Genesee Scientific | 25-500 |
| GenClone Fetal Bovine Serum (FBS) | Genesee Scientific | 25-514 |
| Penicillin-Streptomycin 100X Solution | Genesee Scientific | P4333-100ML |
| DPBS, no Calcium, no Magnesium | Thermo Scientific | 14190250 |

| | | |
|---|---|---|
| Trypsin-EDTA, o.25% 1X, phenol red | Genesee Scientific | 25-510 |
| Paraformaldahyde 20% Aqueous Sol. EM Grade | Electron Microscopy Sciences | 15713 |
| BSA Lyophilized Powder, Fraction V | Genesee Scientific | 25-529 |
| Fibrinogen Type 1-S from Bovine Plasma | Sigma-Aldrich | F8630-1G |
| Thrombin from Bovine Plasma | Sigma-Aldrich | T7513-500UN |
| Aprotinin Protease Inhibitor | Thermo Scientific | 78432 |
| Cytodex Microcarrier Beads | Sigma-Aldrich | C3275-10G |
| CellTracker Deep Red | Thermo Scientific | M22426 |
| Dimethyl Sulfoxide (DMSO) | Sigma-Aldrich | D2650-5X10ML |
| Silencer™ Negative Control No. 1 siRNA | Thermo Scientific | AM4611 |
| Sun1 siRNA | Thermo Scientific | siRNA ID: HSS118638 |
| Sun2 siRNA | Thermo Scientific | siRNA ID: s24465 |
| Nesprin 1 siRNA | Horizon Discovery | J-014039-05-0020 |
| Nesprin 2 siRNA | Horizon Discovery | J-019259-09-0020 |
| Nesprin 3 siRNA | Thermo Scientific | siRNA ID: 129486 |
| Lamin-A siRNA | Thermo Scientific | siRNA ID: s8221 |
| NE-PER Nuclear and Cytoplasmic Extraction Reagents | Thermo Scientific | 78833 |

| | | |
|---|---|---|
| GenBuilder™ Cloning Kit | GenScript | L00701 |
| Neon™ Transfection System 100 µL Kit | Thermo Scientific | MPK10025 |
| STEMdiff™ Endothelial Differentiation Kit | Stem Cell Technologies | 8005 |
| STEMdiff™ Mesoderm Induction Medium | Stem Cell Technologies | 5220 |
| mTeSR™ Plus | Stem Cell Technologies | 100-0276 |
| Heparin Solution | Stem Cell Technologies | 7980 |
| Corning Matrigel Matrix | Fischer Scientific | CB-40234A |
| Animal Component-Free Cell Dissociation Kit | Stem Cell Technologies | 5426 |
| CD31 MicroBeads, mouse | Miltenyi Biotech | 130-097-418 |
| CD45 MicroBeads, mouse | Miltenyi Biotech | 130-052-301 |
| Myelin Removal Beads II, human, mouse, rat | Miltenyi Biotech | 130-096-733 |
| LS Columns | Miltenyi Biotech | 130-042-401 |
| Collagenase/Dispase | Sigma | 10269638001 |
| DNAse | Sigma | 10104159001 |
| Normal Donkey Serum, Freeze-dried | Jackson Immunoresearch | 017-000-121 |
| Click-iT™ EdU Cell Proliferation Kit for Imaging, Alexa Fluor™ 488 dye | Thermo Scientific | C10337 |

| | | |
|---|---|---|
| EZ-Link™ Sulfo-NHS-Biotin | Thermo Scientific | 21217 |
| SuperSignal™ West Femto Maximum Sensitivity Substrate | Thermo Scientific | 34094 |
| BCA Protein Assay Kit | Thermo Scientific | 23225 |
| cOmplete™, Mini, EDTA-free Protease Inhibitor Cocktail | Sigma | 11836170001 |
| PhosSTOP™ | Sigma | 4906845001 |

**Supplementary Table 2. Drugs.**

| **Name** | **Vendor or Source** | **Catalog No./ Clone** | **Working Concentration** |
|---|---|---|---|
| Five1 | MedChem Express | HY-124757 | 2 µM |
| Nocodazole | MedChem Express | HY-13520 | 1 nM |
| Latrunculin A | MedChem Express | HY-16929 | 0.1 µM |
| Lonafarnib | MedChem Express | HY-15136 | 10 µM |
| Progerinin | MedChem Express | HY-159834 | 5 µM |

**Supplementary Table 3. Antibodies.**

| **Target Antigen** | **Vendor or Source** | **Catalog No./ Clone** | **Working Concentration** |
|---|---|---|---|
| Nup98 Antibody (C-5) Alexa Fluor® 647 | Santa Cruz Biotechnology | sc-74578 AF647 | 1:1000 (IHC) |
| UNC84A (Sun1) Polyclonal Antibody | Thermo Scientific | PA5-52548 | 1:500 (IHC)<br>1:1000 (WB) |

| | | | |
|---|---|---|---|
| Lamin A Monoclonal Antibody (133A2) | Thermo Scientific | MA1-06101 | 1:500 (IHC) |
| Anti-gamma H2A.X (phospho S139) antibody | Abcam | ab11174 | 1:500 (WB) |
| Phospho-Histone H2A.X (Ser139) (20E3) Rabbit mAb #9718 | 9718S | Cell Signaling | 1:1000 (WB) |
| NUP98 (C39A3) Rabbit mAb #2598 | Cell Signalling | 2598S | 1:1000 (WB) |
| Nesprin 1 Monoclonal Antibody (MANNES1A(7A12)) | Thermo Scientific | MA5-18077 | 1:1000 (WB) |
| Nesprin 2 Monoclonal Antibody (K20-478-5) | Thermo Scientific | MA5-18075 | 1:1000 (WB) |
| Nesprin 3 Polyclonal Antibody | Proteintech | 27132-1-AP | 1:1000 (WB) |
| Anti-GFP Antibody (Rabbit) | Abcam | Ab290 | 1:1000 (WB) |
| NF-kappaB p65 (D14E12) Rabbit Monoclonal Antibody | Cell Signaling | 8242 | 1:1000 (WB) |
| Phospho-NF-κB p65 (Ser536) (93H1) Rabbit mAb | Cell Signaling | 3033S | 1:1000 (WB) |
| Vimentin (D21H3) Rabbit Monoclonal Antibody | Cell Signaling | 5741 | 1:1000 (WB) |
| Phospho-Vimentin (Ser56) Antibody | Cell Signaling | 3877 | 1:1000 (WB) |
| eNOS Antibody | Cell Signaling | 9572 | 1:1000 (WB) |

| | | | |
|---|---|---|---|
| Phospho-eNOS (Ser1177) Antibody | Cell Signaling | 9571 | 1:1000 (WB) |
| HIF-1 alpha Antibody | Cell Signaling | 3716 | 1:1000 (WB) |
| $\alpha/\beta$-Tubulin Antibody | Cell Signaling | 2148 | 1:1000 (WB) |
| $\beta$-Actin Antibody | Cell Signaling | 4967 | 1:1000 (WB) |
| Focal Adhesion Kinase (FAK) Antibody | Cell Signaling | 3285 | 1:1000 (WB) |
| Phosphorylated FAK (Tyr576) Antibody | Cell Signaling | 3281 | 1:1000 (WB) |
| Cofilin Antibody | Cell Signaling | 5175 | 1:1000 (WB) |
| Phosphorylated Cofilin (Ser3) Antibody | Cell Signaling | 3311 | 1:1000 (WB) |
| GM130 (D6B1) Rabbit Monoclonal Antibody (Alexa Fluor® 647 Conjugate) | Cell Signaling | 59890S | 1: 1000 (IHC) |
| Human VE-Cadherin Antibody | R&D | AF938 | 1:500 (IHC) |
| Streptavidin, Alexa Fluor™ 647 Conjugate | Thermo Scientific | S21374 | 1:1000 (IHC) |
| Alpha-Smooth Muscle Actin Monoclonal Antibody (1A4), eFluor™ 570, eBioscience™ | Thermo Scientific | 41-9760-82 | 1:500 (IHC) |
| Alpha-Smooth Muscle Actin Monoclonal Antibody (1A4), Alexa Fluor™ 488, eBioscience™ | Thermo Scientific | 53-9760-82 | 1:500 (IHC) |

| | | | |
|---|---|---|---|
| Mouse Podocalyxin Antibody | R&D | MAB1556 | 1:500 (IHC) |
| BD Pharmingen™ Alexa Fluor® 647 Rat Anti-Mouse CD144 | BD Biosciences | 562242 | 1:500 (IHC) |
| Anti-Vimentin Antibody (Mouse Aorta) | Abcam | ab92547 | 1:1000 (IHC) |
| ERG Alexa Fluor 647 | Abcam | AB196149 | 1:500 (IHC) |
| Isolectin GS-IB4 From Griffonia simplicifolia, Alexa Fluor™ 488 Conjugate | Thermo Scientific | I21411 | 1:1000 (IHC) |
| Isolectin GS-IB4 From Griffonia simplicifolia, Alexa Fluor™ 568 Conjugate | Thermo Scientific | I21412 | 1:1000 (IHC) |
| Alexa Fluor™ 488 Phalloidin | Thermo Scientific | A12379 | 1 uM (1:200) |
| Alexa Fluor™ 647 Phalloidin | Thermo Scientific | A22287 | 1 uM (1:200) |
| Alexa Fluor™ 555 Phalloidin | Thermo Scientific | A34055 | 1 uM (1:200) |
| Goat anti-Rabbit IgG (H+L) Secondary Antibody, Alexa Fluor 488 | Thermo Scientific | A11008 | 1ug/mL (1:500) |
| Donkey anti-Rabbit IgG (H+L) Secondary Antibody, Alexa Fluor 555 | Thermo Scientific | A31572 | 1ug/mL (1:500) |
| Donkey anti-goat IgG (H+L) Secondary Antibody, Alexa Flour 488 | Thermo Scientific | A11055 | 1ug/mL (1:500) |

| | | | |
|---|---|---|---|
| Donkey anti-Goat IgG (H+L) Cross-Adsorbed Secondary Antibody, Alexa Fluor 555 | Thermo Scientific | A21432 | 1ug/mL (1:500) |
| Goat Anti-Rabbit IgG Antibody (H+L), Peroxidase (PI-1000-1) | Vector Laboratories | PI-1000-1 | 1:1000 |
| Horse Anti-Mouse IgG Antibody (H+L), Peroxidase | Vector Laboratories | PI-2000-1 | 1:1000 |

**Supplementary Table 4. Plasmids.**

| **Name** | **Vendor or Source** | **Catalog No.** |
|---|---|---|
| psPAX2 | Addgene | 12260 |
| VSVG | Addgene | 8454 |
| mCherry-Nesprin-1αKASH (dominant negative Nesprin) | Addgene | 187015 |
| pKLV2-EF1a-NucGFP (nuclear localized control) | Addgene | 184696 |
| pCDH_blast_MCS_Nard_GFP_D50 (Progerin) | Addgene | 167339 |
| pCDH_blast_MCS_Nard_GFP_LAMIN (full length Lamin-A) | Addgene | 167340 |
| mCherry-Vimentin-7 | Addgene | 55156 |
| eGFP-Sun1 | Addgene | 213515 |
